# Childhood exploration drives population-level innovation in cultural evolution

**DOI:** 10.1101/2025.03.28.645934

**Authors:** Elena Miu, Marc Malmdorf Andersen, Sheina Lew-Levy, Felix Riede

## Abstract

The societal effects of children’s learning in cultural evolution have been underexplored. Here, we investigate using agent-based models how a propensity for early exploration in childhood contributes to cultural adaptation and the evolution of long human childhood. Using a complex cultural task, we implemented a two-stage strategy for exploring this space – children explore broadly, more likely to learn new behaviours, while adults exploit behaviours already known, incrementally improving them. We found that populations that followed this two-stage strategy achieved higher payoffs in the long term than populations using the two exploration strategies in a random order. Our models point at a ‘just right’ length of childhood – neither too long, nor too short – allowing individuals enough time to explore before exploiting what they learned. Social learning increased payoffs when agents could copy individuals of a variety of ages. Payoffs decreased under environmental change, especially for long childhoods, because adults did not have enough time to recover between bouts of change.

## Introduction

Culturally learned abilities are at the heart of what makes *Homo sapiens* successful as a species. Through cultural evolution—iterative, trans-generational social transmission and innovation—humans have been able to design and refine new cultural traits which have helped us adapt to diverse environments (1–3). Most cultural evolutionary research has focused on how cultural traits are transmitted and maintained in populations over generations. The mechanism by which novel traits originate remains less well understood.

Here, we define ‘innovation’ as novel traits in a population. When cultural evolutionary scientists have asked questions about innovation they have mainly been working from a population-level perspective: previous work shows that larger groups might generate more innovation (4,5), that network structure is important for collective problem-solving by maintaining more diversity (6–8), and that copying and learning must be used flexibly both individually and collectively in order to produce salient innovations at the population level (9,10). Mechanistically, this work tends to treat innovation either as a random event akin to biological mutations (11–13), or a result of recombination (14,15).

Less attention has been paid to the individual characteristics that drive the emergence of truly novel cultural traits within populations (16). Existing research suggests that certain personality types or cognitive propensities may make some individuals more likely to innovate than others (17–20). Importantly, these individual contributions may not arise purely from random variation but instead reflect more stable, systematic patterns across populations. Over longer temporal scales, evidence from developmental psychology and anthropology suggests that childhood, as a distinct life stage, consistently contributes to innovation across diverse contexts. Drawing on these insights, we here investigate whether children, through a unique combination of cognitive characteristics, are likely to be drivers of innovation at the individual and the population level.

The role of children in cultural evolution has so far been underplayed. There has been a tendency to focus on the receptive and copying abilities of children more so than their potential for innovation across longer periods of time (see (21) for review). When considered, children have been seen as the vehicle for high-fidelity intergenerational transmission – culture is only useful if maintained faithfully from generation to generation (22–24). Given that children need to learn this cultural knowledge, it has been assumed that children are psychologically predisposed for faithful learning and imitation (25–27).

Children have been portrayed as less proficient than adults at generating adaptive innovations, and more prone than adults to make copying errors (28–31). Studies of cultural macroevolution have similarly focused on transmission errors (13,32) and on when the capacity for and mechanisms of faithful cultural transmission developed (33–35) rather than how material culture variation and innovation came about.

Recent work in developmental psychology contests some of these assumptions. Experimental research suggests that children are highly exploratory in their learning and are intrinsically motivated to pay costs for novel information, whereas adults are more conservative, specializing and relying more on previously acquired knowledge and skills (36–39). This contrast is partly due to differences in cognitive processing: children often operate with what in Bayesian terms is referred to as ‘flat priors’, meaning they are less constrained by existing assumptions and more open to exploring a wider range of possibilities (40). Adults’ stronger priors, built on accumulated experience, can lead to cognitive rigidity and a preference for exploiting known strategies over exploring novel ones. Sherratt and Morand-Ferron (41) provide an additional explanation for adults’ conservatism, arguing that the diminishing future value of novel information with age discourages exploration. Older individuals tend to know more, making novel information rarer and often less useful. Furthermore, the limited time remaining in their lifetimes to exploit new discoveries reduces the payoff of exploration, reinforcing their preference for established knowledge.

Children’s exploratory behaviour not only fuels their own learning but also makes them a critical source of variation within populations. Play, a hallmark of childhood, is characterized by exploration, experimentation, and variation (42), providing a mechanism for children to introduce novel behaviours and ideas into their environments. In play, children are intrinsically motivated by the rewards provided by reducing uncertainty and acquiring new information, which makes exploration and information gathering both enjoyable and effective for them (43,44). This intrinsic drive to seek out and resolve uncertainty positions children as agents of cultural variation, as they naturally generate novel hypotheses and test them through play and other exploratory behaviours (45,46).

Complementary observations from within the field of anthropology further suggest that a prolonged costly childhood may have evolved as a period for individuals to acquire the wealth of cultural knowledge on which humans rely for survival (47–49). Play is one of the mechanisms through which this learning takes place (50). Play is not only a way to rote-learn relevant physical and social behaviours (51); in diverse societies, children have been shown to be active agents who guide their own learning, often in vicarious and unsupervised ways through exploratory play (52–54). Such play provides low-cost and low-risk opportunities to explore and potentially discover new behaviourial variants, including those that involve material culture, another hallmark domain of human behavior (55). Moreover, children are often provided with functional minitures – toys – of the technologies they will encounter later in life, which allows them to explore material and mechanical affordances in the context of exploratory play (56) and unconstrained by time and energy burdens of reproduction and care. Considering this literature, Riede et al. (57) have argued that the sweet spot for innovation lies prior to adulthood, specifically in late childhood and adolesence.

Gopnik (58) has interpreted the differences between adults and children to be that they do not lie at different ends of an exploratory-conservative spectrum, but that they actually use contrasting strategies. This split-strategy life history may represent an evolved solution to the explore-exploit problem all organisms face: when to learn novel information and when to use the information they already possess. Previous work on the explore-exploit tradeoff shows that some of the ways organisms have adapted to solve this issue include division of labour and diversification among individuals (59). Gopnik’s work instead proposes that differential exploration within the same individuals is also a solution, and thus, that childhood is a period evolved specifically for exploratory learning.

Importantly, children are often raised in environments that differ substantially from those their parents experienced, exposing them to new tools, technologies, and social contexts. This generational shift in environmental conditions enables children to develop skillsets and knowledge that diverge from those of their parents, equipping them with unique perspectives and approaches to problem-solving. These differences in experiences and learning opportunities allow children to adapt to novel circumstances and, in turn, introduce variation into their cultural and social groups. By the time children become adults, they possess distinct abilities and insights shaped by their exploratory learning and exposure to different environments. This makes children a critical source of cultural variation, with implications for innovation, individual development, and the broader adaptive dynamics of cultural evolution.

While children’s exploratory nature might promote innovation in adulthood within the same individuals, innovation is not just an individual-level process. Population-level innovation is an incremental process through which individuals build on other’s solutions (10,60). Adult-like exploitation of known behaviours is beneficial under environmental stability (61), especially in worlds where cultural adaptation is prevalent and powerful (62). In contrast, adult specialisation loses relevance if the environment changes because specialised knowledge is no longer up-to-date with current environmental conditions. Further, the time spent investing in specialisation is lost. It has thus been suggested that broader repertoires of solutions are more useful under elevated environmental variation – both in the human evolutionary past and more recently (63–65). Research suggests that children can be a source of variation and diversity that popualtions can build on. For example, modelling studies show that learning from younger individuals can act as a middle ground between social learning and innovation, and thus, can represent an adaptive strategy during periods of environmental fluctuation (66,67). Similarly, Lew-Levy and Amir (21) review empirical cases in which children’s innovations helped communities adapt to social and ecological change.

It seems, then, that children’s flexibility, play, and their exploratory nature should make them ideal sources of variation and, later on, innovation, which could have important consequences for the evolution of cultural adaptation at the population level. Here, we aim to understand the conditions under which such a life history of learning could have evolved, and their consequences for innovation at the population-level. To do so, we integrate children’s exploration and learning into a framework designed to study the evolution of cultural adaptation and cumulative cultural evolution. We explore how childhood exploration relates to (i) innovation later in life and (ii) innovation outcomes at the population-level. We then (iii) investigate how this changes with environmental variation. While previous work has often pitted individual and social learning against each other (61,62,68,69), the aim of this study is primarily to understand how different types of individual learning are leveraged across life history to find the best solutions, and to explore the consequences of a mix of types of individual learning and social learning at the population level.

We designed an agent-based model in which agents explored a problem space designed to capture key features of complex problems that organisms face (Fig. 1a). The agents’ goal was to find the best payoff of a set of cultural options or traits, each associated with its own payoff. Conceptually, these can be thought of as different types of behaviours relevant to survival – different foraging techniques, or the skill neded to make and use different sorts of tools. Each trait was associated with a number of skill levels, or expertise levels, such that increasing levels were associated with increasingly higher payoffs. Each increasing level can be thought of as a more complex, or more effective version of the lower levels. In this way, we are capturing one type of dependency characteristic to cultural systems (70). We designed a steep problem space, where many solutions were ineffective, and only few solutions were effective, to model the kinds of difficult problems that (human) groups have been able to solve historically through cultural adaptations. Each turn, the environment changed with probability *pEnv*. Change involved drawing a new set of base payoffs and their associated increments.

**Fig. 1.**
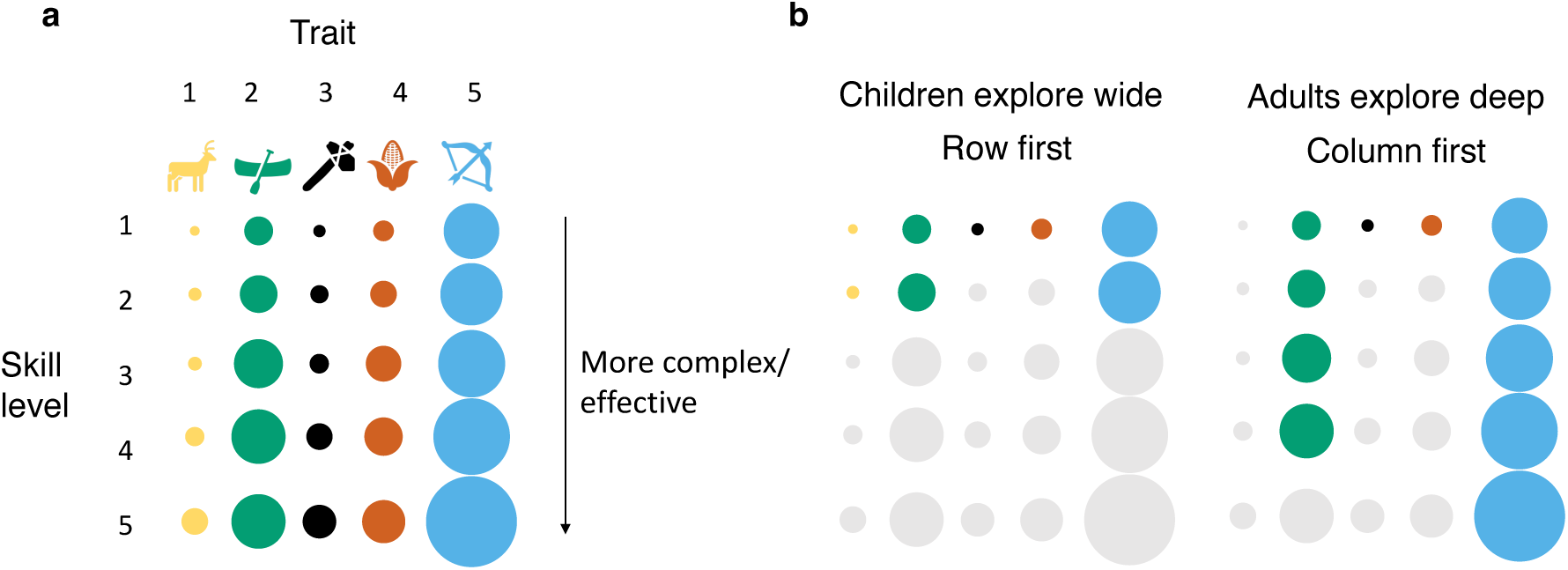
Problem space and exploration. (a) Problem space consisting of a number of traits, each associated with its own payoff. Each trait can be improved by increasing its level, each increase associated with increasing payoffs relative to the base level. (b) Two individual learning strategies, for each age stage of the agent. Children explore widely, while adults explore deeply.

We distinguished between two life history stages, each associated with a different way of exploring the problem space (Fig. 1b). In keeping with the evidence from the literature, when the agents were children they explored the space more widely, i.e. they were more likely to learn the easier versions of the traits but finding it harder to learn the higher levels – they *explored*. Adults, in contrast, explored the space depth-wise, being more likely to specialise and improve what they already knew, but not learn new traits at the lower levels – they *exploited*.

Agents were born naïve and searched the problem space by learning a behaviour they did not know each step. We implemented two individual search and learning strategies: *explore*, in which an agent picked a level 1 behaviour it did not know yet, and learned its accurate payoff; and *exploit*, in which an agent picked the best behaviour it knew already, and increased its skill level, thus learning the newly improved behaviour and its associated increased accurate payoff. The agents had two life stages. When they were children, they *explored* (as defined above), and when they are adults they *exploited*. Agents aged from childhood to adulthood at a set threshold we call *length of childhood*.

To assess whether agents implementing this two-stage learning strategy were successful at exploring the space, we compared a population of two-stage agents with a control population of agents who used both *explore* and *exploit* in a random order (as opposed to sequentially, as agents with childhood do), but in the same proportion as the agents with childhood. Thus, if the proportion of the childhood was 20% of the lifetime, then the control agents would use *explore* 20% of the time – the probability of *explore* over the lifetime *pExplore* was therefore also the proportion of childhood in our model.

The agents could also learn new behaviours from each other through social learning. We implemented two versions of social learning: 1. Random social learning, where agents picked a random model in the population, picked a random behaviour that model knows, and learned it and its payoff; 2. Best (i.e. payoff-biased) social learning, where agents picked a random model from the population and from its repertoire learned the behaviour with the highest payoff. Both types of social learning allowed agents to skip ahead and learn behaviours or skill levels they did not already know. In order to keep the number of learning opportunities constant within each *lifetime*, social learning was traded off with individual learning according to a parameter *pS*, which we varied. Thus, *pS* dictated the probability of learning through social learning instead of individual learning (i.e. *explore* or *exploit*) each turn. The probability of learning through individual learning was, therefore, *1-pS* each turn.

We implemented an overlapping generations population structure. On each turn a given agent could die with a set probability *pDie*, in which case the agent was removed from the population and replaced with a naïve agent. We varied *pDie*, which was defined as 1/*lifetime*, such that on average each agent in this condition would live an average of *lifetime* rounds. In this model, we were only concerned with what the agents knew, not how they used this information to reap payoffs (i.e. production costs), therefore the measure of a successful individual was the highest payoff they new so far.

## Results

### Lifetime and *pExplore*

Payoffs were maximised for agents with long lifetimes and at intermediate *pExplore* for both the control condition and the childhood condition (Fig. 2a-b). A longer lifetime meant agents had more learning opportunities, which translated into an increased chance to discover the best payoffs. For *pExplore*, however, the relationship with highest payoff was non-linear: when *pExplore* was small, the agents did not learn enough traits at skill level 1 in order to find a good option to invest in – in this case, agents started exploiting too early to reach optimal payoffs. Similarly, when *pExplore* was high, agents spent too much time exploring at the detriment of exploitation opportunities (note that exploitation very quickly gave rise to much higher payoffs than exploration; see Figs. S1, S2). The highest payoffs were achieved at intermediate proportions of childhood (i.e. *pExplore* = 0.2), where the opportunities for exploration were balanced out most optimally with exploitation.

**Fig. 2.**
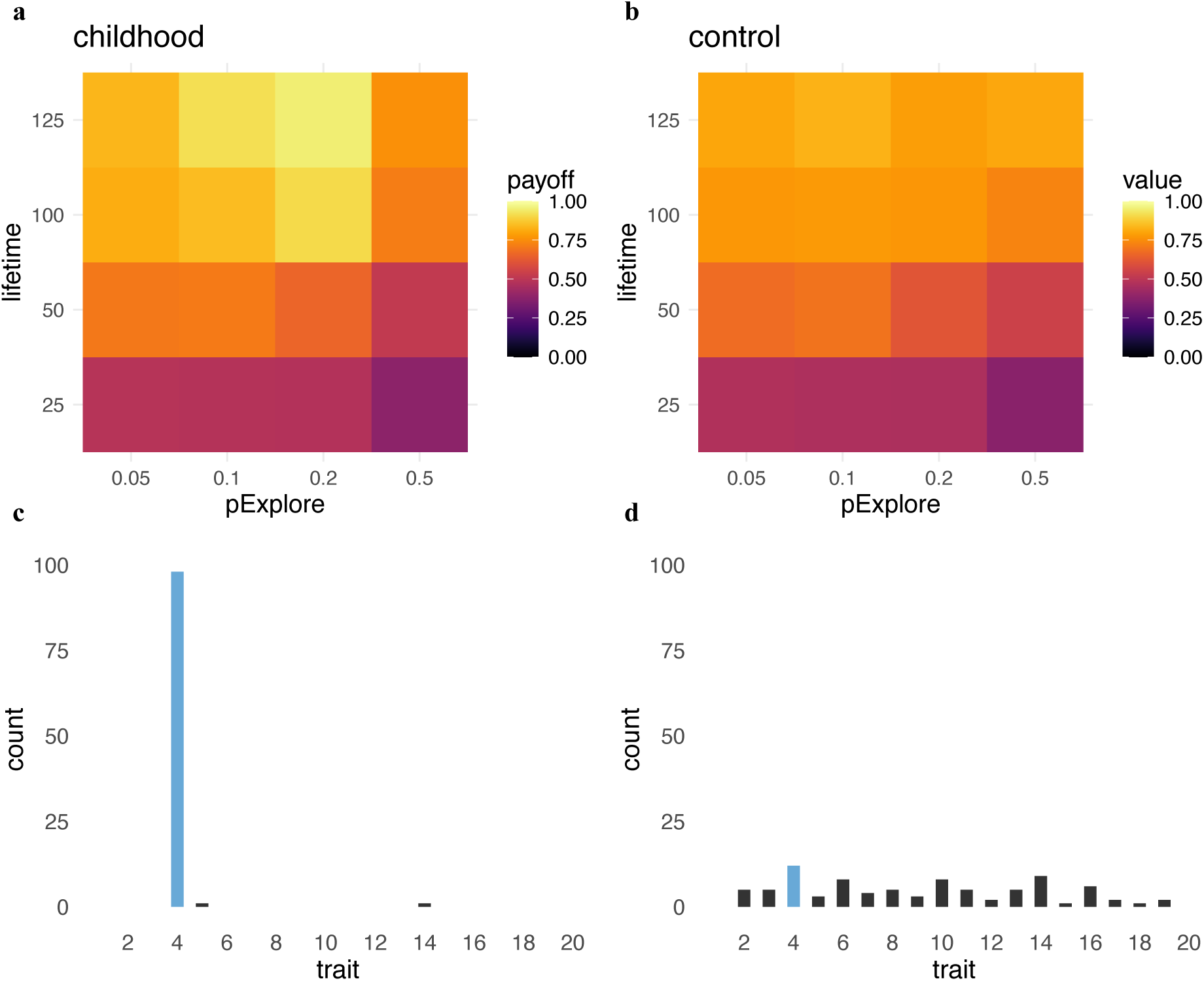
Payoffs for varying lengths of lifetime and probabilities of *explore*. Median values of maximum payoffs for varying lifetime lengths and varying *pExplore* (i.e. probability agents use explore/proportion of childhood) for both (a) the childhood and (b) the control conditions. Median values calculated over 100 agents, 20 repeat simulations, and over the last generation of a five-generation run, with *pS* = 0, *pEnv* = 0. Distributions of which traits individual agents exploit, for agents in (c) the childhood and (d) the control condition. Results from one run of the simulation, with *lifetime* = 100, *pExplore* = 0.2, *pS* = 0, *pEnv* = 0. In this run, option 4 (coloured in blue) was the option that resulted in the highest payoff.

The benefit of childhood relative to control (i.e. the difference between the childhood condition and the control condition) was also maximised at high lifetimes and intermediate *pExplore* values (Fig. 2a-b). Importantly, as the agents in both conditions used the same number of *explore* rounds and the same number of *exploit* rounds, this difference was purely due to the order in which they use these. This period of initial exploration allowed most agents with childhood to find and exploit the option best matched to their environment, i.e. the one that would eventually lead to maximum payoff compared to agents in the control condition who, as a group, exploited a variety of options (Fig. 2c-d). For shorter lifetimes, we found no difference between the childhood and control conditions (Fig. 2a-b, *lifetime* 25 and 50).

### Social learning

Random social learning increased payoffs in overlapping populations for both the childhood and the control condition, but only to a point (Fig. 3). The higher *pS*, the lower the proportion of exploration, as exploration and social learning also trade off in our model. A population needs enough useful information to begin with, which then can spread (69). When *lifetime* was high, higher social learning drove the population to almost maximum payoffs (Fig. 3, S5, S6), but when the *lifetime* was lower improvement was slower (Fig. S3, S4). These patterns held for all values of *pExplore* (Figs. S3-6).

**Fig. 3.**
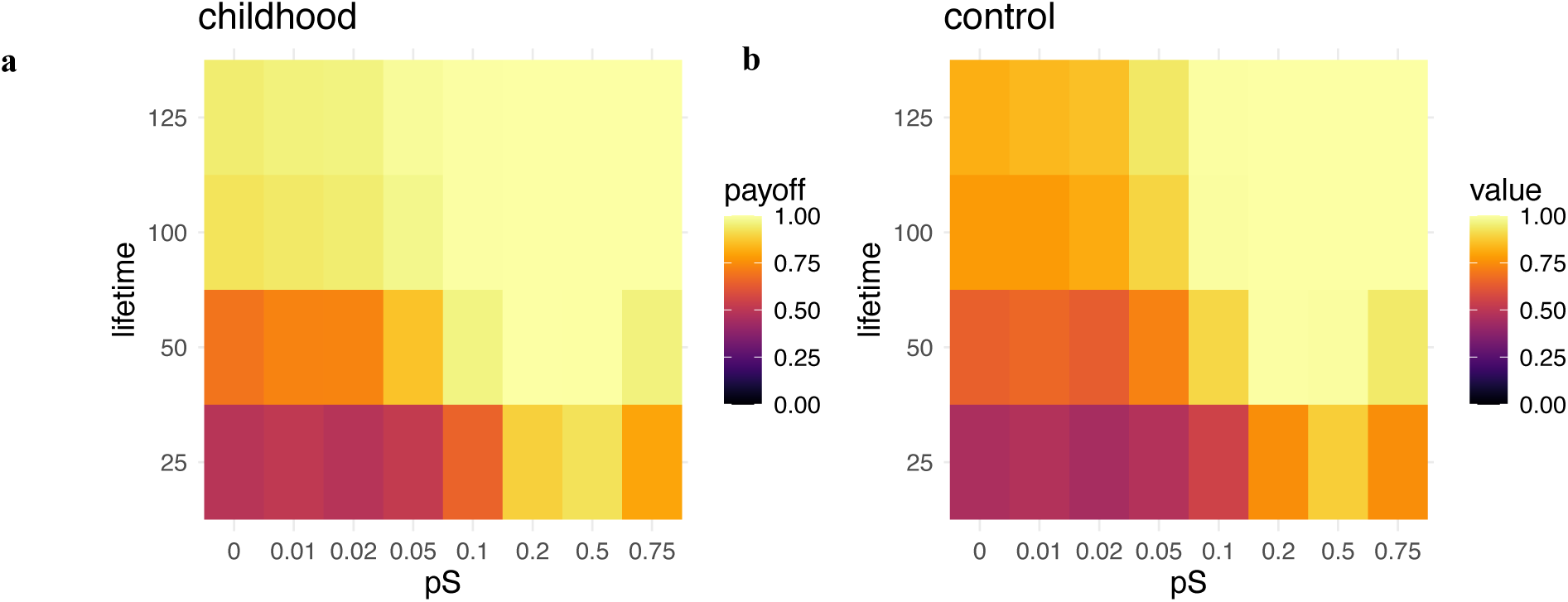
Payoffs for varying probabilites of social learning. Median values of maximum payofs for varying probabilities of social learning *pS* (when social learning was random) and varying *lifetimes*, for (a) childhood and (b) control. Medians calculated over 100 agents, 20 repeat simulations, and over the last generation of a 5-generation run, with *pExplore* = 0.2, *pEnv* = 0.

**Fig. 4.**
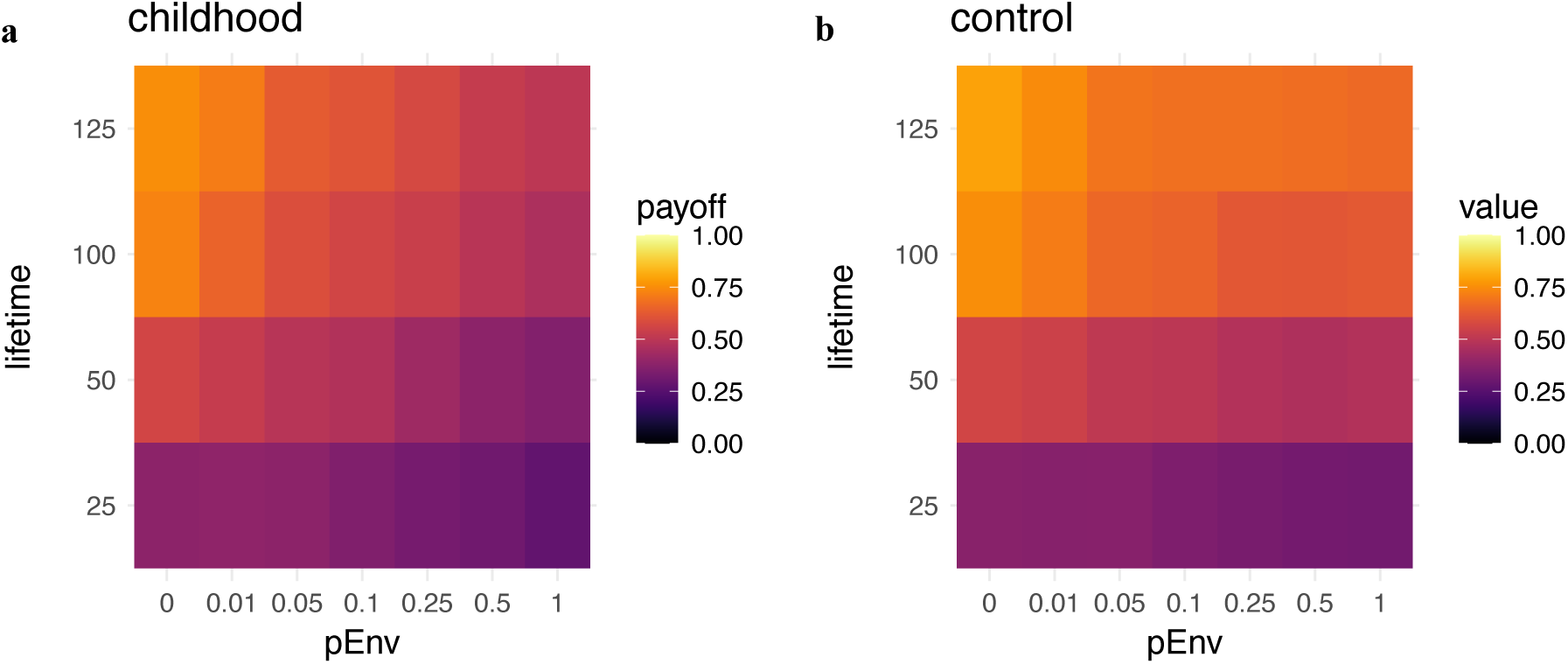
Payoffs for varying probabilities of environmental change. Median values of maximum payoffs for varying probabilities of environmental change *pEnv* and varying *lifetimes,* for both (a) childhood and (b) control. Medians calculated over 100 agents, 20 repeat simulations, and over the last generation of a five-generation run, with *pExplore* = 0.5, *pS*=0.

**Fig. 5.**
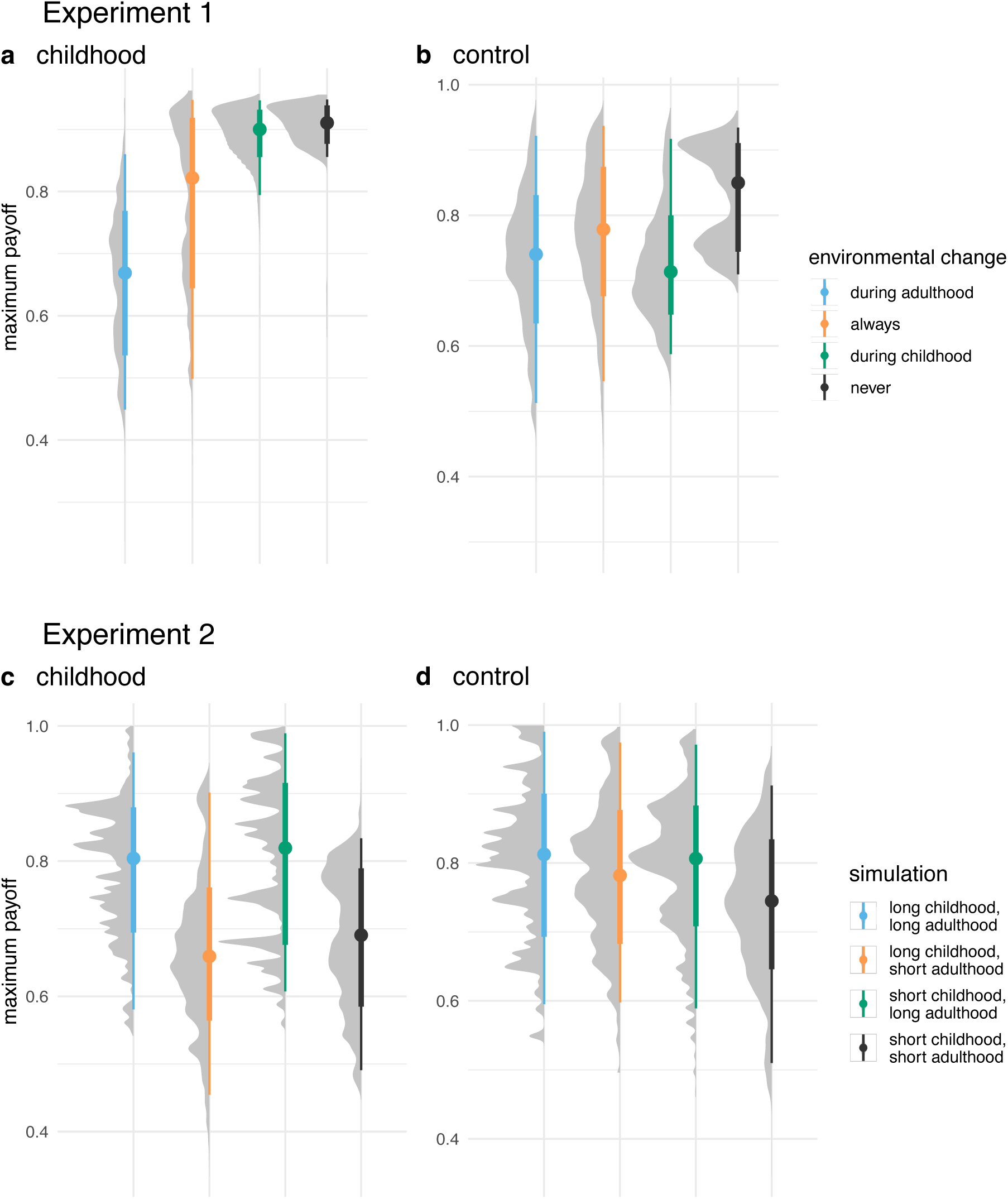
Results from environmental change experiments. Distributions of maximum payoffs for simulations manipulating when environmental change takes place, for (a) childhood and (b) control, using *lifetime* = 100, *pExplore* = 0.5, *pEnv* = 0.2. Distributions of maximum payoff from simulations systematically manipulating the lengths of childhood and adulthood, for (c) childhood and (d) control, using *pExplore* = 0.5, *pEnv* = 0.2. Results from 10 repeat simulations each. The points indicate the median of the distribution, the thick line indicates the interquartile ranges, and the shape indicates the shape of the density distribution.

**Fig. 6.**
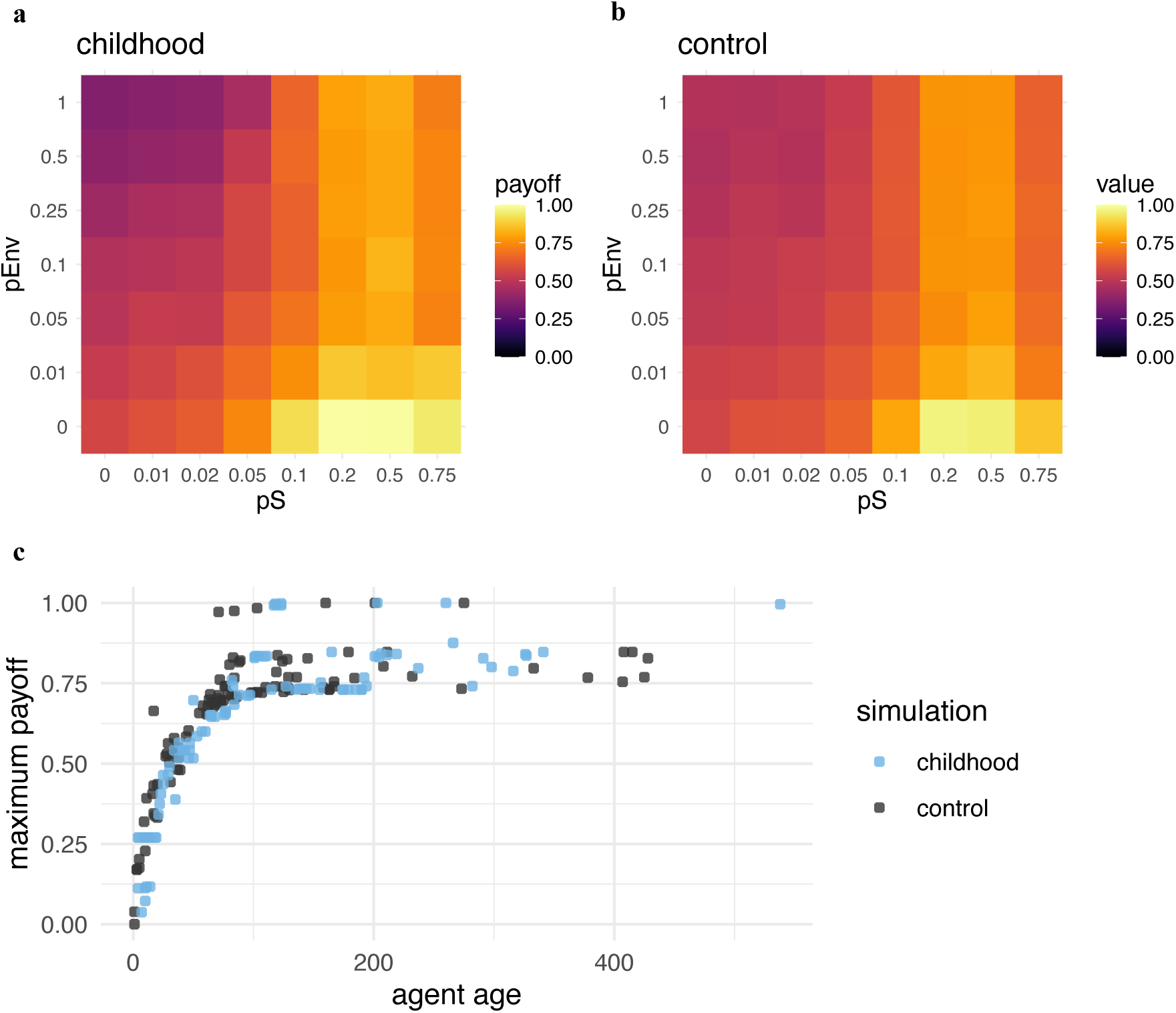
Payoffs for varying combinations of probabilities of social learning and environmental change. (a-b) Median values of maximum payoffs for varying probabilities of social learning pS (when social learning was random) and varying probabilities of environmental change *pEnv,* for both conditions. Distributions over 100 agents, 20 repeat simulations, and over the last generation of a 5-generation run, with *lifetime* = 50, *pExplore* = 0.5. (c) Maximum payoffs as a function of agent age. Results from one run of a simulation with *N* = 100 agents, *lifetime* = 100, *pExplore* = 0.2, *pS*=0, and *pEnv* = 0.2. Snapshot from round 555, after an environmental change took place.

Childhood populations achieved higher payoffs than control populations at different *pS* values that varied by *lifetime* and were moderated by *pExplore*. Adding social learning revealed more conditions under which childhood was useful: when lifetime was high, childhood was more useful than control under low *pS,* but increasing *pS* quickly removed the differences between conditions (Fig. 4, S5, S6), but when lifetime was short childhood conditions achieved higher payoffs than control conditions at intermediate and higher values of *pS*. This was especially true for higher *pExplore* (Figs. S3, S4). This suggests that the benefit of chilhood relative to control was highest under intermediate social learning, when the option best matched to the environment could spread more quickly (Fig. 3), before the control condition could catch up.

When social learning was payoff-biased (i.e. agents copied the best option of their randomly chosen model), these patterns still held, but at an accelerated pace (Figs. S7-10). For lifetimes over 50, the populations discovered the option with the highest payoff in the entire grid and converged on it. The differences between the childhood and control conditions vanished very quickly under payoff-biased social learning.

### Environmental change

Environmental change reduced payoffs (Fig. 4, Figs. S11-14). As expected, this was true for longer lifetimes as this means exposure to more environmental change. However, payoffs also decreased when *pExplore* was high, and affected the childhood condition more strongly, for all lifetimes. In the results presented so far, the payoffs in the childhood condition never fell below the payoffs in the control condition, but under high environmental change that was no longer the case – for all values of *lifetime*, the payoffs for childhood were lower than the payoffs for control when p*Env* = 1.

To understand why longer childhoods independent of long lifetimes suffered more under environmental change, and to distentagle whether it is longer childhoods or shorter adulthoods that were responsible for the decrease in payoffs, we performed two experiments within the simulation framework. We fixed agent life and ran one set of simulations where agents were born and died at the same time to see how environmental change affected individuals.1) We manipulated when environmental change took place (during adulthood, childhood, both, or never) in both the childhood and the control condition (with *lifetime* = 100, *pExplore* = 0.5, *pEnv* = 0.2); 2) We systematically manipulated the lengths of childhood and adulthood under environmental change in both conditions, such that agents had either a short childhood (50 rounds) and a short adulthood (50 rounds), a short childhood (50 rounds) and a long adulthood (100 rounds), a long childhood (100 rounds) and a short adulthood (50 rounds), or a long childhood (100 rounds) and a long adulthood (100 rounds). For this we used *pExplore* = 0.5, *pEnv* = 0.2.

We found that for the control condition, the timing of environmental change did not matter – whether it took place during childhood, adulthood, or both, it reduced payoffs to the same amount (Fig. 5b, experiment 1). For the childhood condition, however, the timing was crucial – if environmental change took place during childhood, this did not decrease the payoffs at all, while if it took place during adulthood it lowered the payoffs the most (Fig. 5a). When we systematically manipulated the lengths of childhood and adulthood (experiment 2), we found that for the childhood condition it was not the length of the total lifetime that mattered, but the length of adulthood (Fig. 5c). Long adulthoods achieved higher payoffs than short adulthoods, while the length of childhood did not matter. In the control condition, payoffs did not depend on the length of childhood or adulthood (Fig. 5d), but on lifetime and therefore exclusively on the number of learning opportunities.

In the childhood condition, therefore, payoffs did not suffer under childhood, and a long childhood did not increase payoffs compared to a short childhood. However, payoffs did suffer under adulthood, but a longer adulthood also increased payoffs compared to a shorter one. This explains why populations with long childhoods suffered more under high *pExplore* – the proportion of time spent in adulthood was shorter, meaning they had less time to recover after an environmental change event.

### Recovering after environmental change

We were interested in whether social learning could help populations recover from decreases in payoff caused by environmental change. We found that payoffs did indeed increase even under high environmental change conditions when random social learning was used (Fig. 6a-b, Figs. S14-S17). For example, when *pExplore* = 0.5, the condition under which environmental change was most detrimental, populations with *lifetime* = 50 experienced an increase of almost 100% in payoffs even when the environment changed every turn (i.e. *pEnv* = 1) when using random social learning 20% of the time (Fig. 6a). As before, social learning under these conditions removed the differences between the childhood and the control conditions. Payoff-biased social learning showed similar patterns, with higher payoffs achieved at high *pS* (Figs. S18-S21). These patterns held for all values of *lifetime* and *pExplore*.

Older agents generally knew better options but payoff was non-linearly related to agent age, as expected from the diminishing returns shape of the payoff increment with payoffs tapering off around age 100 (Fig. 6c). The highest payoffs were not necessarily known by the oldest individuals – here, after an environmental change event, younger, but not the youngest, adults knew the behaviours with the highest payoffs (Fig. 6c).

The results presented here were robust under different increment shapes, when we relaxed the assumption that exploitation acts on the best option, and when we used alternative controls (see Supplementary Text).

## Discussion

We developed an agent-based model to assess the benefit of a split explore early-exploit later strategy under a variety of social learning and environmental change conditions. We found that the benefit of exploratory childhood was highest during long lifetimes and intermediate lengths of childhoods, a result consistent with the evolved constellation of human childhood (71). Social learning increased payoffs, but only in populations with overlapping age structures (72), where agents could copy individuals of a variety of ages. However, it was not always the oldest individuals who held the best knowledge. The benefit of chilhood was highest under intermediate social learning, when the optimal option could spread quicker than agents learning randomly could catch up. Finally, payoffs decreased under environmental change. This was not because agents suffered during childhood, suggesting that childhood was indeed a protected period, but because these agents did not have enough time during adulthood to catch up on their payoffs through exploitation. This suggests that under conditions of harsh environmental change, a shorter childhood could be beneficial, as documented in observational studies (73–76).

With social learning, however, even under severe environmental change populations could recover. Our work integrates life history and cultural evolution (66,77) and adds to a growing literature incorporating specific individual cognitive mechanisms in frameworks designed to study cultural transmission processes in order to understand population-level patterns of behaviour, especially those that focus on the role of children (78–80).

It might seem curious that the strategy we chose as our control performed so well, especially under social learning and environmental change conditions, but previous work has shown that strategies that incorporate variation in sophisticated ways tend to be successful in a wide range of conditions (14,81). Our task models a type of cumulative cultural evolution where new niches are discovered early in life, and then optimised by adults through focused competency (10). This conceptualisation allows for the type of path dependence we see in real world cultural traits, where initial choices constrain future ones (70,82,83). This can prove problematic under the type of random environmental change we implemented in our model, where the path taken might become suboptimal after a shift. Through the randomness inherent in the control condition, individuals and populations using this strategy could maintain more path diversity, and shift to a more effective path after an environmental change, while the agents in our childhood condition were trapped on the path they specialised in. Notably, extended childhoods are the exception rather than the norm in nature.

We found that early exploration leads to better solutions both within the same individuals later in life, and for other individuals in the population. It has been suggested that exploratory play is an important driver of both learning and innovation, a type of ontogenetic niche construction wherein, through manipulating the appropriate type of materials – often miniature adult tools (84) – children familiarise themselves with the physical affordances and social context of tool use (85–87). This familiarity with the material is believed to enable innovation later in life when the children grow up to make their own tools, and recent work has linked this initial period of exploratory material play with adaptation to environmental change (88): Meyer and Riede present data on the differential survival of two Greenlandic groups throughout the Little Ice Age in the fourteenth century CE – Inuit foragers who thrived, and Norse farmers who did not. The authors show that Inuit toy repertoires were rich and diverse, while the Norse toy repertoires were characterised by few toys, mosly related to highly normative adult roles as part of the prevalent farming lifestyle. They propose that the diversity of the Inuit toy repertoire is not just a proxy for the diversity of the adult tool repertoire that might have helped the Inuit during a period of climactic shifts, but also that playing with a diverse repertoire, alongside a famously encouraging Inuit pedagogy, primed children for an innovative mindset, that would bestow them the flexibility to invent and shift to new tools during hardship. Here, we add to this and show that exploration is not just beneficial to the same individuals later in life, but to other individuals in the population as they are searching for better solutions (Figs. 5 and 7).

Highlighting the role of exploration and self-guided play for learning does not detract from the importance of teaching for both socialisation and cultural evolution (26,34,89–91). Teaching, either formal or informal, is a large part of how children learn the social and norms as well as the functional skills needed to survive in their natal society. Teaching is also considered necessary for the kind of cumulative cultural evolution that our species is particularly successful at, where innovations are added to information that needs to be accurately passed on between generations (92–94). How faithfully this information needs to be reproduced is a matter of domain – some have suggested that more functionally restricted domains might require more accurate transmission, and therefore more teaching (95), although recent empirical work has found that teaching is more frequent in normative domains and less likely to be mentioned for subsistence and manifacturing skills (96).

Teaching per se, however, is in fact anathema to innovation, and formal instructional relations such as apprenticeship are often more about knowledge control than about fostering creativity (97). Teaching is necessary for cumulative cultural evolution, but how individuals introduce novelty to the information they have learned must be explained by a variation-generation mechanism. Play and exploration are such mechanisms.

If, as shown by our model, play and exploration during childhood lead to better, more innovative solutions, we can make predictions about how this propagates at the group level, and therefore about what group characteristics could promote innovation. Societies that provide children with more agency and space for innovation should, all things being equal, generate more innovation (86). There is evidence that group organisation and group structure can be manipulated to produce more innovative solutions (6,98,99), but to our knowledge there is currently little work, either modelling or observational, that links the propensity of childhood exploration to population-level innovation. Recent work has shown that good ideas spread more successfully when seeded in smaller minority groups that can act as incubators, as opposed to seeded in larger groups (100). Children’s peer cultures (21) can be thought of as the original, evolved population incubators in which children’s developmental propensity for exploration, variation generation, and innovation is realised.

Our results regarding the success of the control condition during environmental change suggest that exploration should not just be restricted to childhood, and does benefit adults as well. In a similar model investigating the evolution of phenotipic plasticity in the context of trade-offs between sampling (i.e. exploring) and specialising, Frankenhuis and Panchanathan (74) found that the greatest reliance on sampling evolved when the environment was fairly uncertain and sampling cues were intermediately reliable, consistent with our results showing that childhood is useful at intermediate social learning and environmental change. They also found that individual differences in plasticity can evolve as a result of their sampling history and how reliable the cues they sample were early on in life. In our model, all agents age at the same time, but future work should investigate the effect of individual differences in exploration propensity and the length of the period of exploration. It is also worth noting that childhood itself is internally differentiated; while very small children may engage in a great deal of exploration, this is unlikely to be efficacious due to cognitive and physical ontogenetic constraints. In teenage just prior to the onset of reproduction, such constraints are lifted but neophily peaks suggesting that it is this period in particular during which salient innovations could appear (101,102). Future models should seek to further differentiate the age structure within multi-generational models, and to even include post-reproductive individuals and their contribution to the knowledge ecology. By the same token, should some adult individuals specialise in the type of playful exploration we conceptualise here, populations could benefit from a type of division of labour that begets specialist innovators who, behaving ‘childlike’, exclusively focus on producing innovations for the population. Some past and contemporary societies have already implemented similar strategies, for instance, through the emergence of dedicated inventors (103), and the implementation of research and educational careers well into reproductive age. Moreover, explicit tinkering pedagogies (104) likewise adopt the stance that playful exploration catalyses innovation independent of age.

Our study has limitations. We focused exclusively on the informational benefits of the early exploration characteristic to children, but childhood is embedded in a complex system of social organisation that include cooperative breeding, wide-range non-kin cooperation, and provisioning for the young. No single factor or approach is likely to satifacorily explain the evolution of the uniquely human childhood (105). Children can afford to explore precisely because adults are providing for them and keeping them safe. While some of the costs and benefits associated with these activities can be subsumed within the payoff structure our model assumes, future work should incorporate these types of costs specifically in order to tease apart the cultural and economic consequences of childhood. Our results are consistent with narrative scenarios of why a grossly extended childhood evolved in the *Homo* linage but allow us to be considerably more specific about the order of selection. Our results suggest, for instance, that childhood developed under stong selection – possibly in the context of variability selection (106,107) – and prior to social learning becoming a major force in cultural evolution. This insight alone allows us to further qualify cultural macroevolutionary scenarios for when and why cumulative culture driven by social learning emerges in hominins (33). More broadly, our novel modeling exercise provides a strong mandate motivating further investigations – observational and experimental – into the role of children in innovation and cultural evolution.

## Materials and Methods

### Task and environment

We assume that within the payoffs of each option are subsumed any costs that might be associated with each type of exploration, and the main costs considered here are represented in the trade-off between sampling the environment and specialisation. For the results presented here, we assumed a problem with *nTraits* = 20 traits and a maximum *sMax* = 100 skill levels.

More specifically, payoffs for the base level (level 1) were drawn from an exponential distribution, and then squared, which resulted in few traits with large payoffs, and many traits with small payoffs (Fig. S1a). Each increasing skill level was associated with a diminishing returns increment, where improvements were initially very beneficial but become less so as the trait became more specialised, to model most skills that develop through incremental improvement and cumulative cultural evolution (10). The increment was the same for all traits. The maximum fully exploited payoffs were on average 20 times higher than the mean basic payoff, to simulate how exploited behaviours are much more effective than behaviours that can be discovered early on.

Each turn, the environment changed with probability *pEnv*. Change involved drawing a new set of base payoffs and their associated increments. The base payoffs were drawn from the same family of distributions as the original payoffs, thus resulting in a different problem of the same kind. After an environmental change, the agents retained their knowledge, and updated the payoffs of their known options according to the new environment.

### Search and learning mechanisms

We use the terms *explore* and *exploit* in keeping with the literature concerning the explore-exploit tradeoff (59,108). In our conceptualisation, *explore* corresponded to individuals discovering new options they did not know already, and *exploit* corresponded to improving and specialising in options they already knew (as opposed to using the option to reap payoff, as it is sometimes used in the literature). All agents had the same goal: to discover better solutions. In our model, exploit was not random – the agent picked the best (i.e. highest payoff) behaviour it knew and exploited it, thus simulating specialisation and reflecting the agents’ aim to find the best solution. To introduce variation, both exploration mechanisms were error-prone: with a certain error probability, both explorers and exploiters would sometimes exploit a random behaviour they already knew.

The length of the lifetime an agent was alive (the number of rounds they explored the space), and the proportion of it spent in childhood were parameters we varied to investigate dynamics arising from the interaction between these key life history parameters (72,109). Note that this is not an evolutionary model – we did not let the parameters evolve but instead we fixed them at different values and compared their effectiveness.

### Population structure

Given the stochastic nature of the death-birth process, some agents did live longer and some shorter lives. Despite variable ages, all agents had the same proportion of childhood within the same population. All populations consist of N=100 agents. We did run auxilliary analyses with N=500 and N=1000 and found very similar results.

We ran each simulation for five generations (i.e. five times the length of lifetime) and collected the maximum payoff of each agent in the last generation. We ran each simulation condition 20 times to average over potential effects due to initial payoff distributions. We standardised payoff values within each simulation repeat run. Because the distributions of maximum payoffs were often skewed, we report median values.

We ran all combinations of the following parameters:

*lifetime* = {25, 50, 100, 125}

*pExplore* = {0.05, 0.1, 0.2, 0.5}

*pS* = {0, 0.01, 0.02, 0.05, 0.1, 0.2, 0.5, 0.75}

*pEnv* = {0, 0.01, 0.05, 0.1, 0.25, 0.5, 1}

*pError* = {0.05}

*N* = {100}

*nTraits* = {20}

*sMax* = {100}

## Data and materials availability

The code for all the analyses is available at: https://osf.io/b26rp/?view_only=2717ec06406c4eaeab9e90a52008e4dc

## Acknowledgments

We thank Tom Morgan for comments on an earlier version of the manuscript.

## Author contributions

Conceptualization: EM, MMA, SLL, FR

Methodology: EM

Investigation: EM

Visualization: EM

Supervision: FR

Funding acquisition: MMA, SLL, FR

Writing—original draft: EM, FR

Writing—review & editing: EM, MMA, SLL, FR

