## Supplementary Materials for "Childhood exploration drives population-level innovation in cultural evolution"

**This PDF file includes:**

Figs. S1 to S23
Supplementary Text

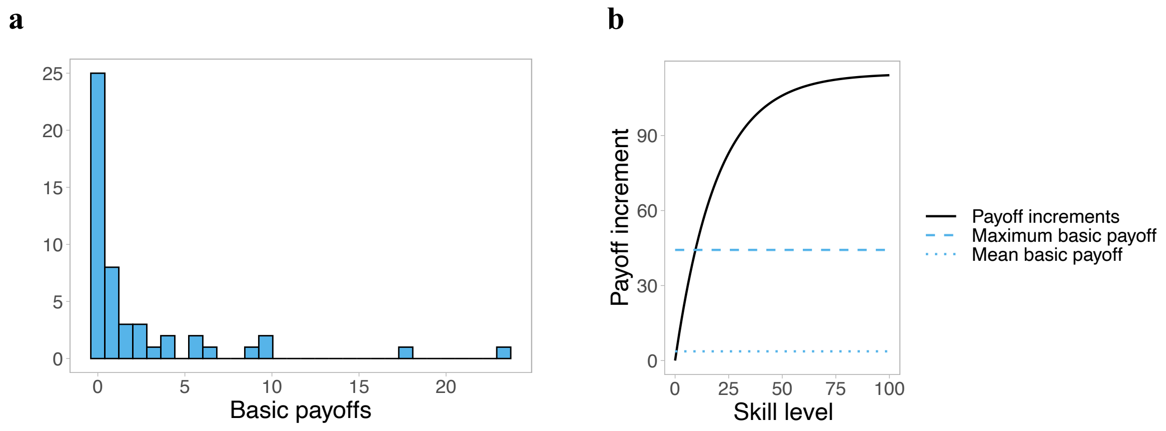


**Fig. S1** – (a) Basic payoff distribution. (b) Payoff increment shape and amplitude relative to base payoffs.

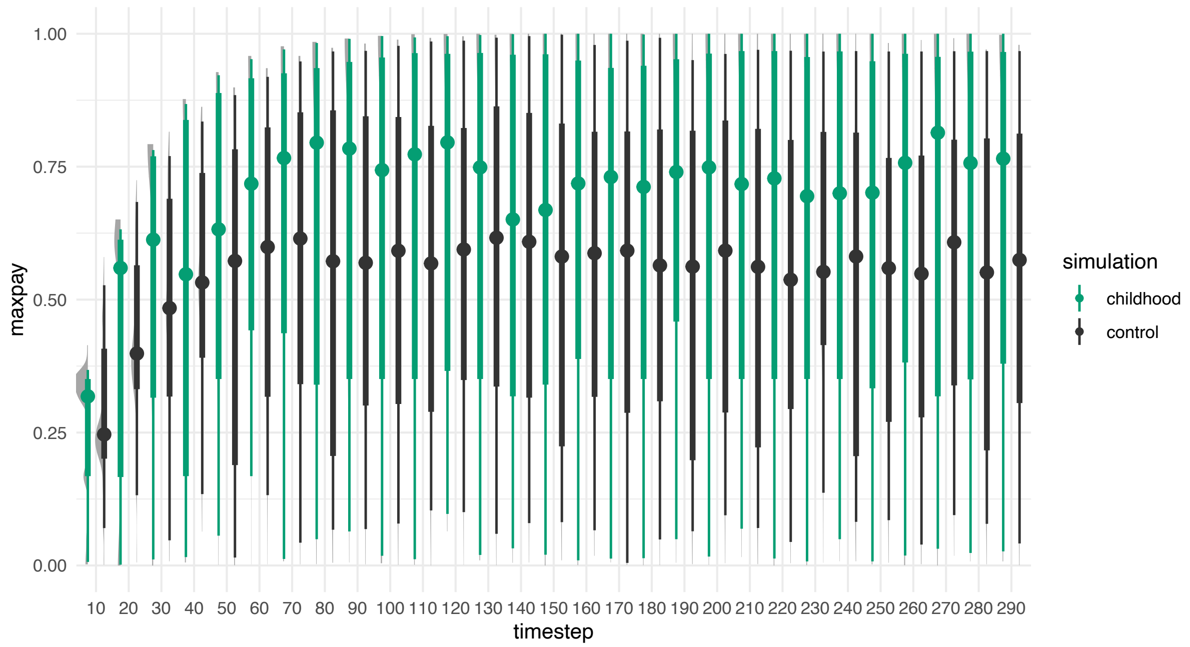


**Fig. S2**– distributions of maximum payoff over time, over 100 agents for the first 300 rounds of a simulation run. Simulation parameters: *lifetime* = 50, *pExplore* = 0.2, *pS* = 0, *pEnv* = 0

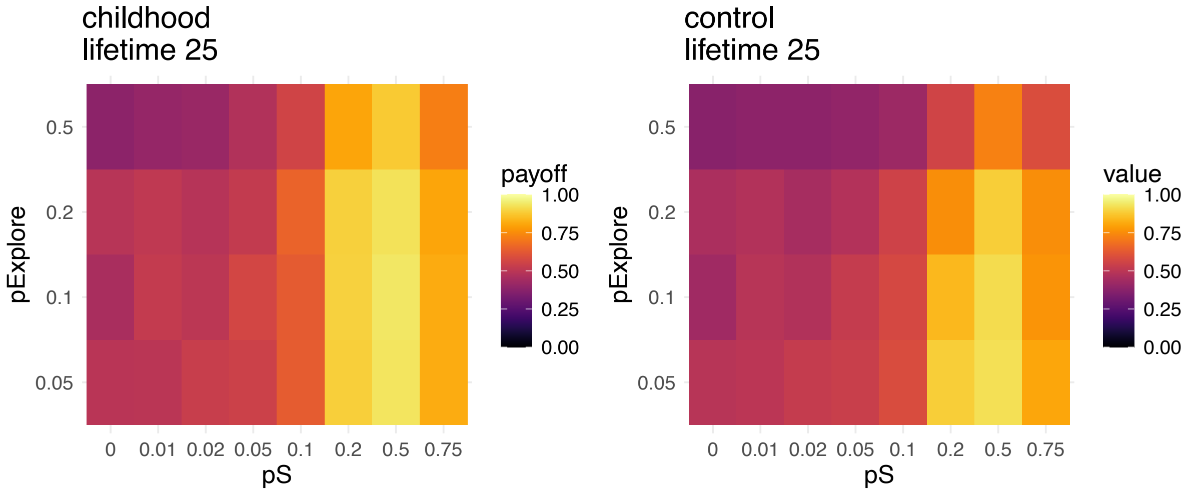


**Fig. S3** – Median values of maximum payoffs for varying probability of social learning *pS* (when social learning is **random**) and varying *pExplore* (i.e. probability agents use explore/proportion of childhood). Medians calculated over 100 agents, 20 repeat simulations, and over the last generation of a 5-generation run, and *lifetime* = 25, *pEnv* = 0.

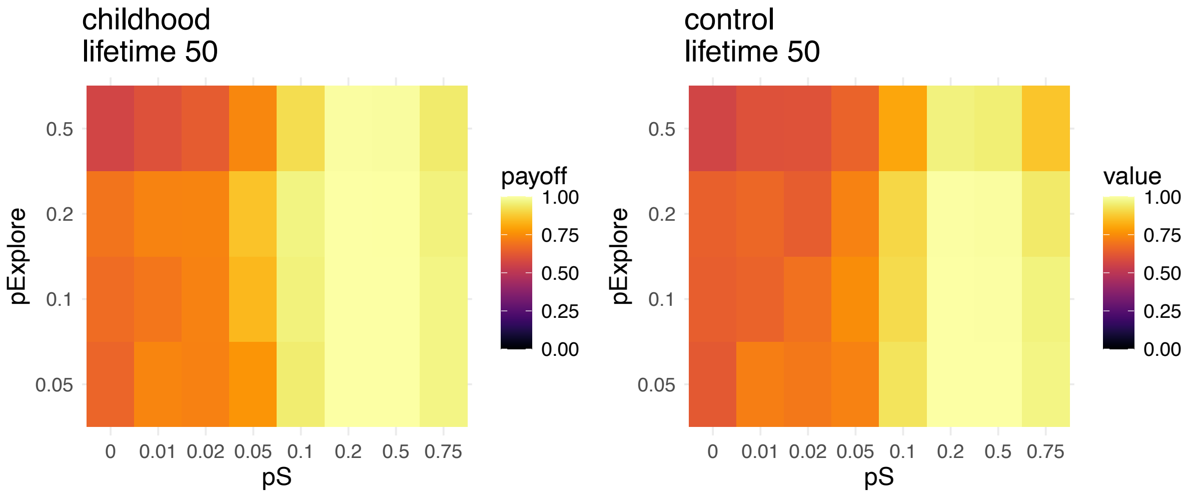


**Fig. S4** – Median values of maximum payoffs for varying probability of social learning *pS* (when social learning is **random**) and varying *pExplore* (i.e. probability agents use explore/proportion of childhood). Medians calculated over 100 agents, 20 repeat simulations, and over the last generation of a 5-generation run, and *lifetime* = 50, *pEnv* = 0.

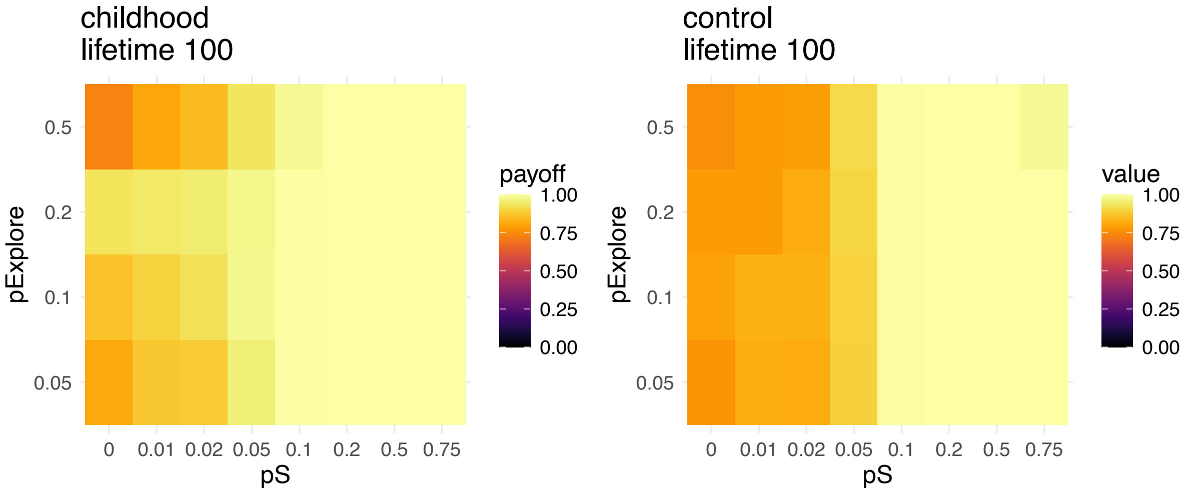


**Fig. S5** – Median values of maximum payoffs for varying probability of social learning *pS* (when social learning is **random**) and varying *pExplore* (i.e. probability agents use explore/proportion of childhood). Medians calculated over 100 agents, 20 repeat simulations, and over the last generation of a 5-generation run, and *lifetime* = 100, *pEnv* = 0.

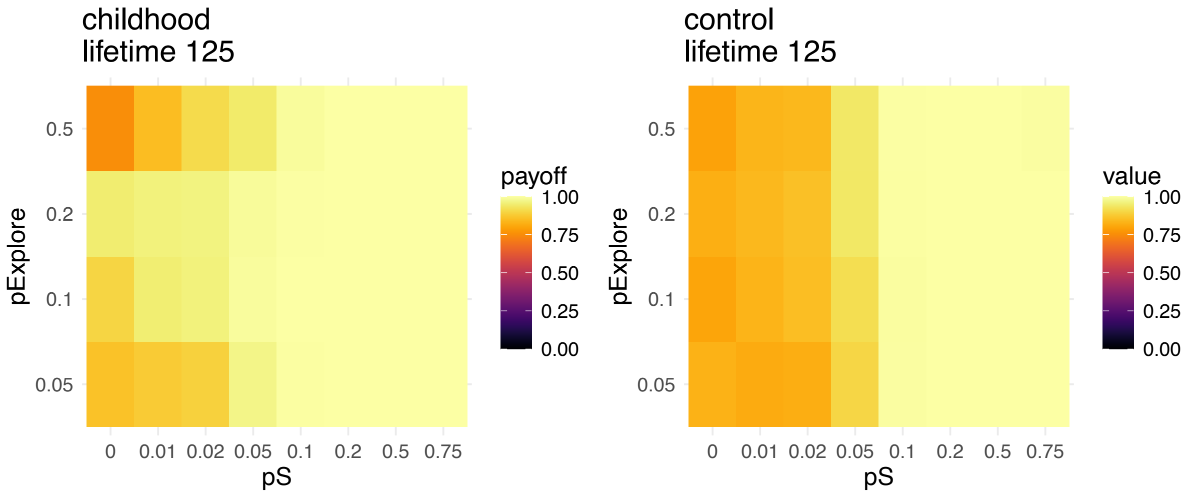


**Fig. S6** – Median values of maximum payoffs varying probability of social learning *pS* (when social learning is **random**) and varying *pExplore* (i.e. probability agents use explore/proportion of childhood). Medians calculated over 100 agents, 20 repeat simulations, and over the last generation of a 5-generation run, and *lifetime* = 125, *pEnv* = 0.

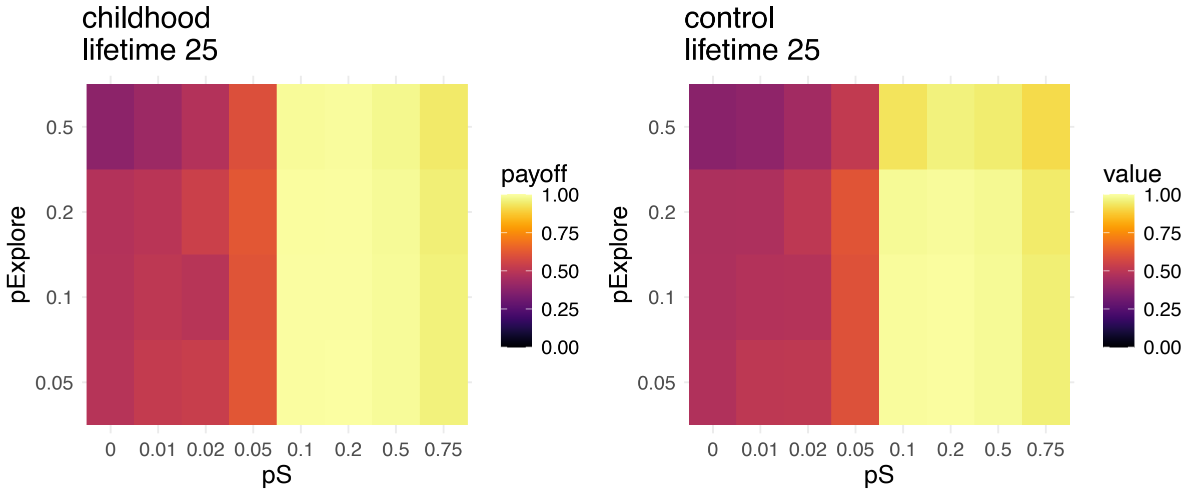


**Fig. S7** – Median values of maximum payoffs for varying probability of social learning *pS* (when social learning is **payoff-biased**) and varying *pExplore* (i.e. probability agents use explore/proportion of childhood). Medians calculated over 100 agents, 20 repeat simulations, and over the last generation of a 5-generation run, and *lifetime* = 25, *pEnv* = 0.

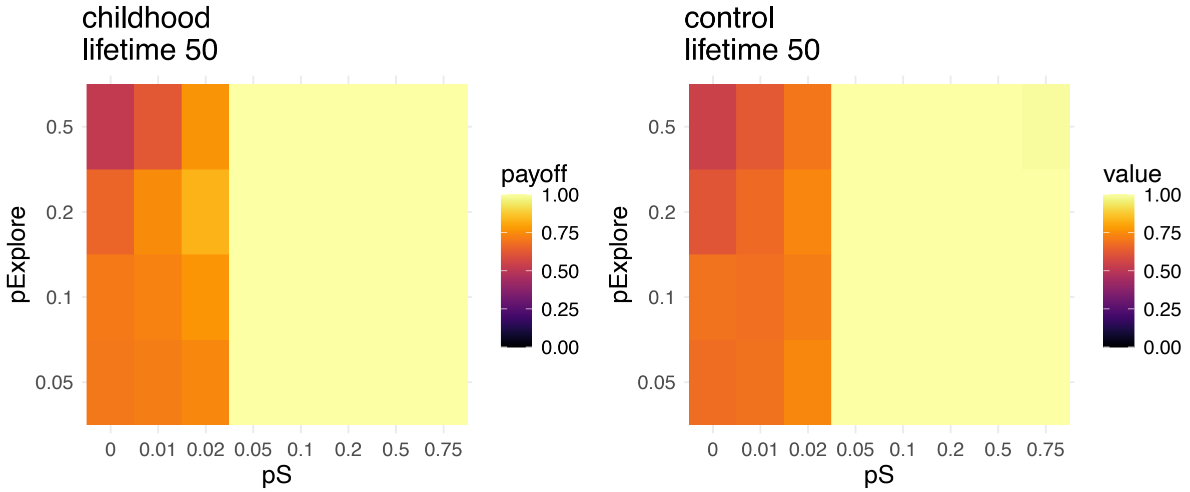


**Fig. S8** – Median values of maximum payoffs for varying probability of social learning *pS* (when social learning is **payoff-biased**) and varying *pExplore* (i.e. probability agents use explore/proportion of childhood). Medians calculated over 100 agents, 20 repeat simulations, and over the last generation of a 5-generation run, and *lifetime* = 100, *pEnv* = 0.

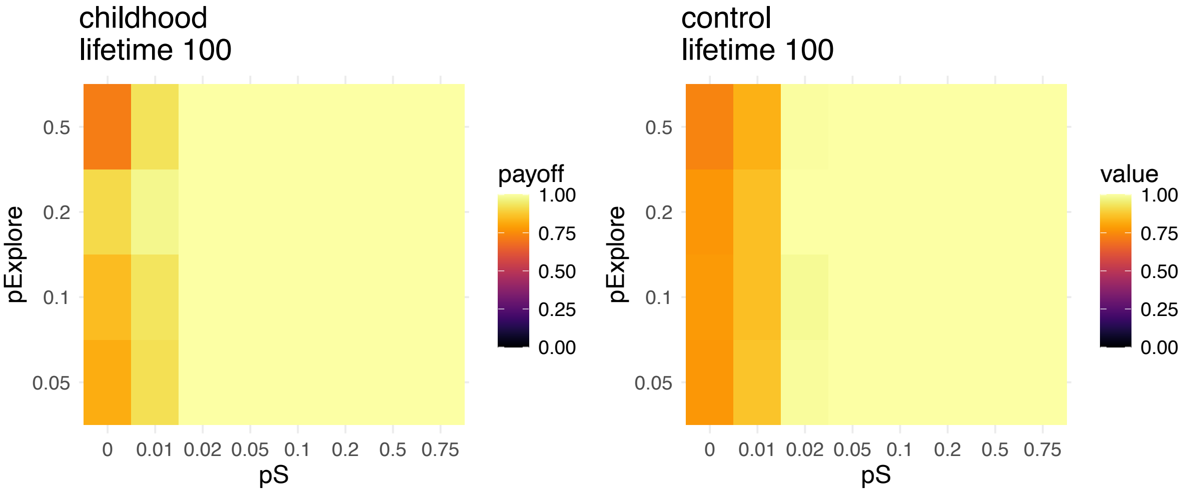


**Fig. S9** – Median values of maximum payoffs for varying probability of social learning *pS* (when social learning is **payoff-biased**) and varying *pExplore* (i.e. probability agents use explore/proportion of childhood). Medians calculated over 100 agents, 20 repeat simulations, and over the last generation of a 5-generation run, and *lifetime* = 100, *pEnv* = 0.
.

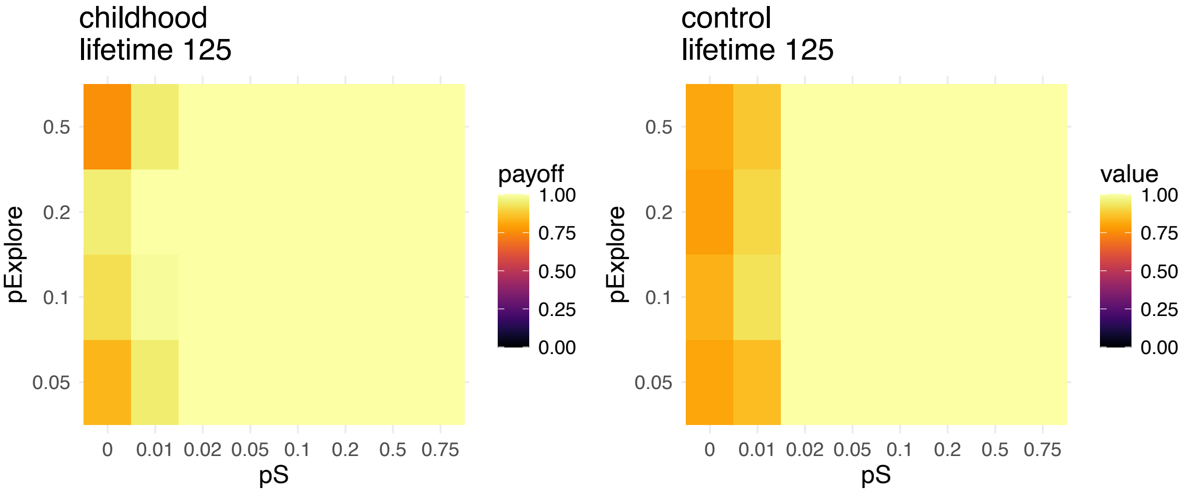


**Fig. S10** – Median values of maximum payoffs for varying probability of social learning *pS* (when social learning is **payoff-biased**) and varying *pExplore* (i.e. probability agents use explore/proportion of childhood). Medians calculated over 100 agents, 20 repeat simulations, and over the last generation of a 5-generation run, and *lifetime* = 125, *pEnv* = 0.

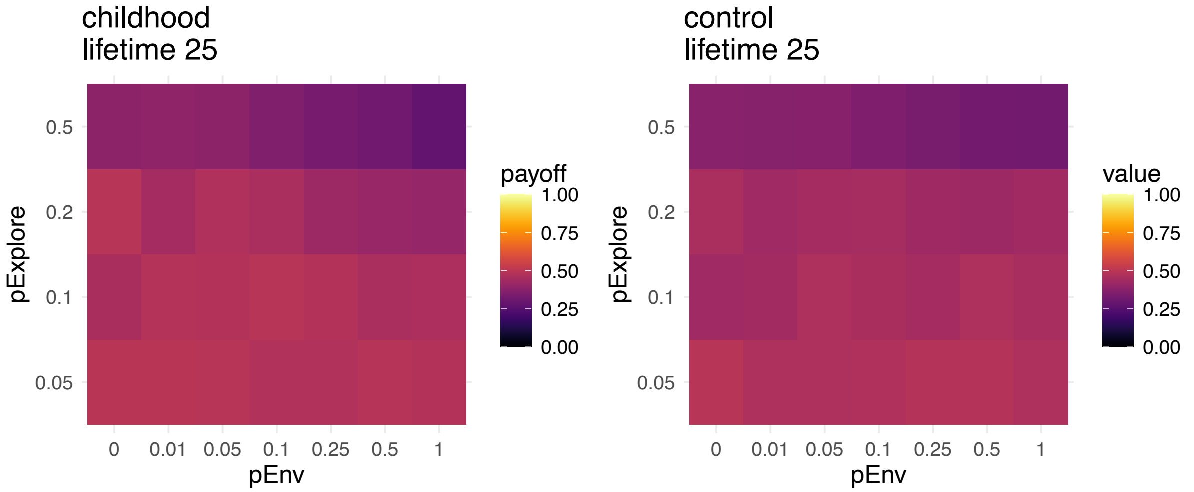


**Fig. S11** – Median values of maximum payoffs for varying probability of environmental change *pEnv* and varying *pExplore* (i.e. probability agents use explore/proportion of childhood). Medians calculated over 100 agents, 20 repeat simulations, and over the last generation of a 5-generation run, and *lifetime* = 25, *pS* = 0.

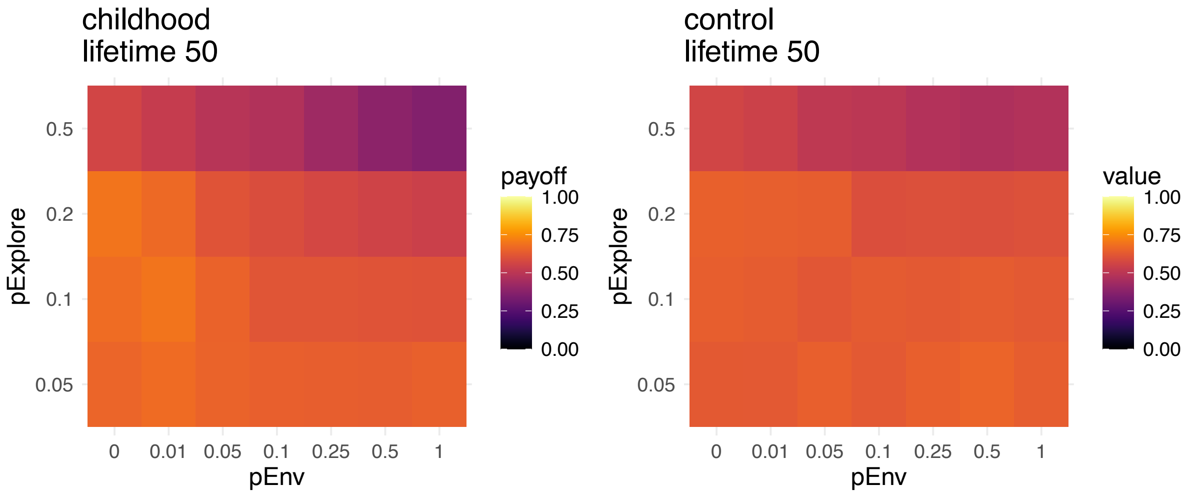


**Fig. S12** – Median values of maximum payoffs for varying probability of environmental change *pEnv* and varying *pExplore* (i.e. probability agents use explore/proportion of childhood). Medians calculated over 100 agents, 20 repeat simulations, and over the last generation of a 5-generation run, and *lifetime* = 50, *pS* = 0.

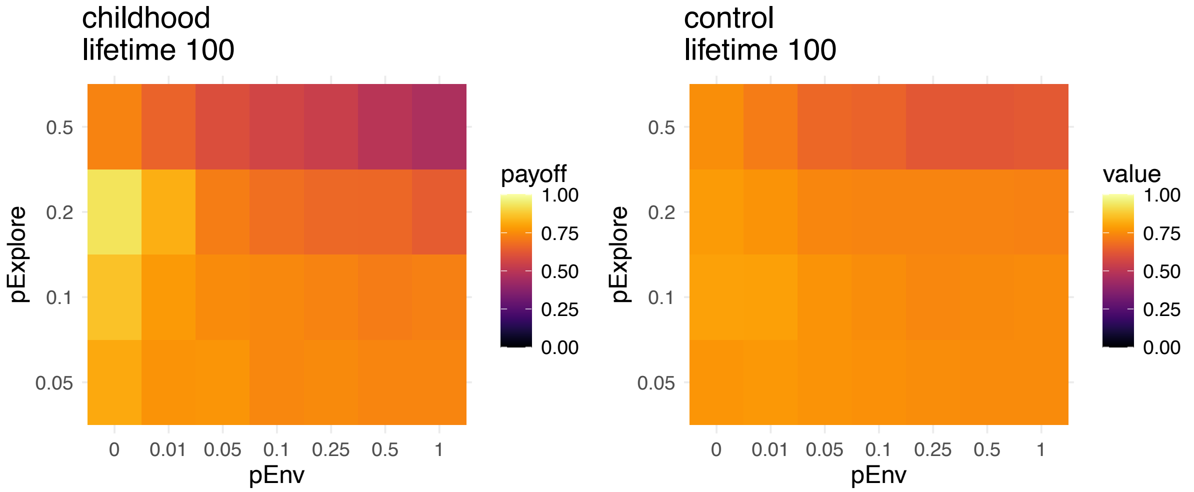


**Fig. S13** – Median values of maximum payoffs for varying probability of environmental change *pEnv* and varying *pExplore* (i.e. probability agents use explore/proportion of childhood). Medians calculated over 100 agents, 20 repeat simulations, and over the last generation of a 5-generation run, and *lifetime* = 100, *pS* = 0.

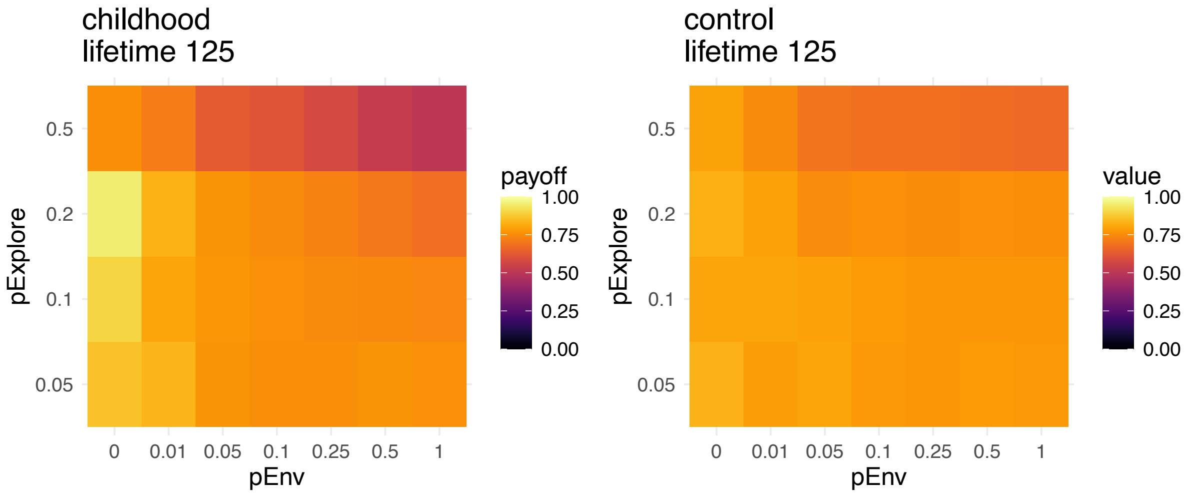


**Fig. S13** – Median values of maximum payoffs for varying probability of environmental change *pEnv* and varying *pExplore* (i.e. probability agents use explore/proportion of childhood). Medians calculated over 100 agents, 20 repeat simulations, and over the last generation of a 5-generation run, and *lifetime* = 125, *pS* = 0.

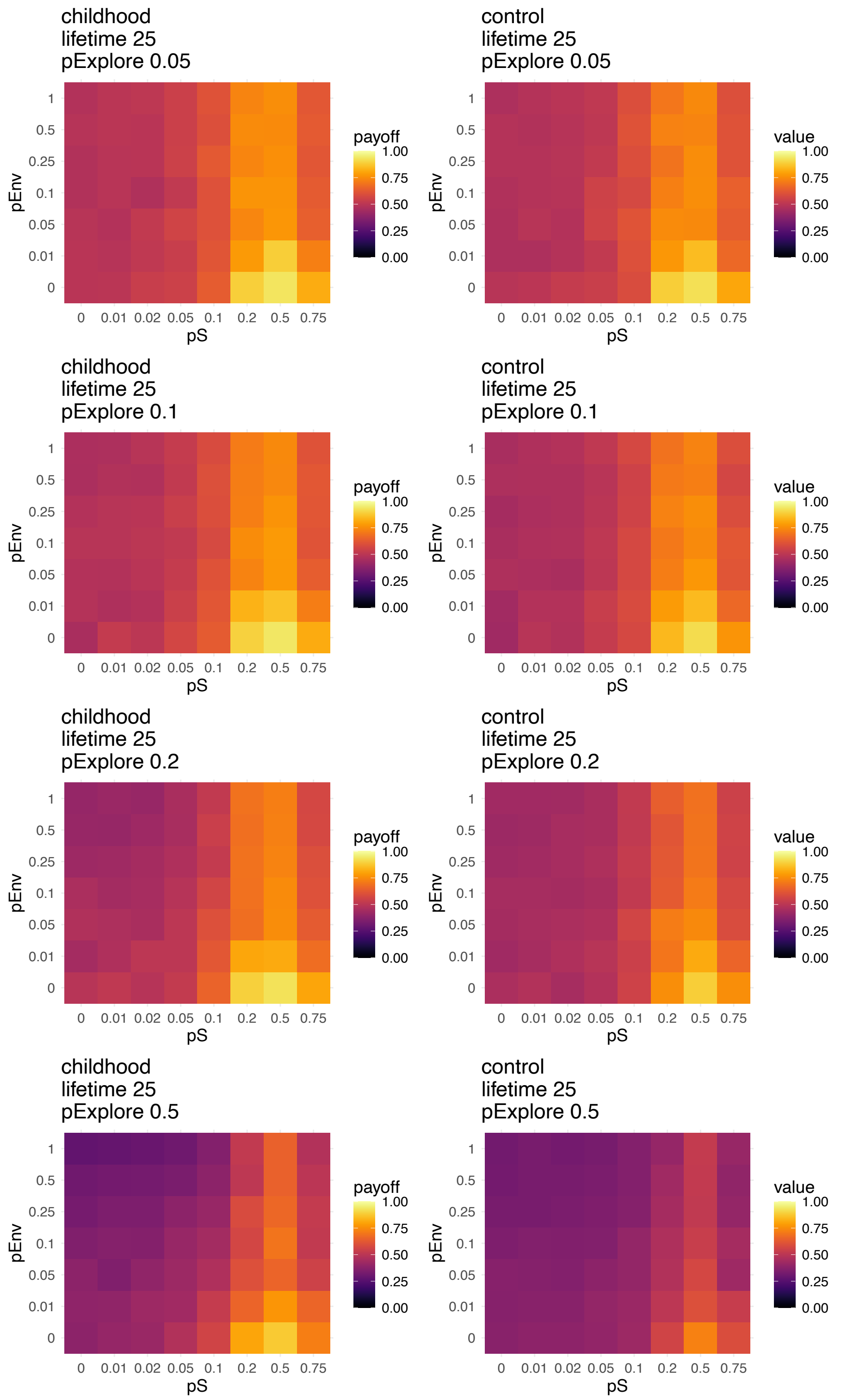


**Fig. S14** – Median values of maximum payoffs varying probabilities of social learning *pS* (when social learning was **random**) and varying probabilities of environmental change *pEnv.* Medians calculated over 100 agents, 20 repeat simulations, and over the last generation of a 5-generation run, and *lifetime* = 25, for all values of *pExplore*.

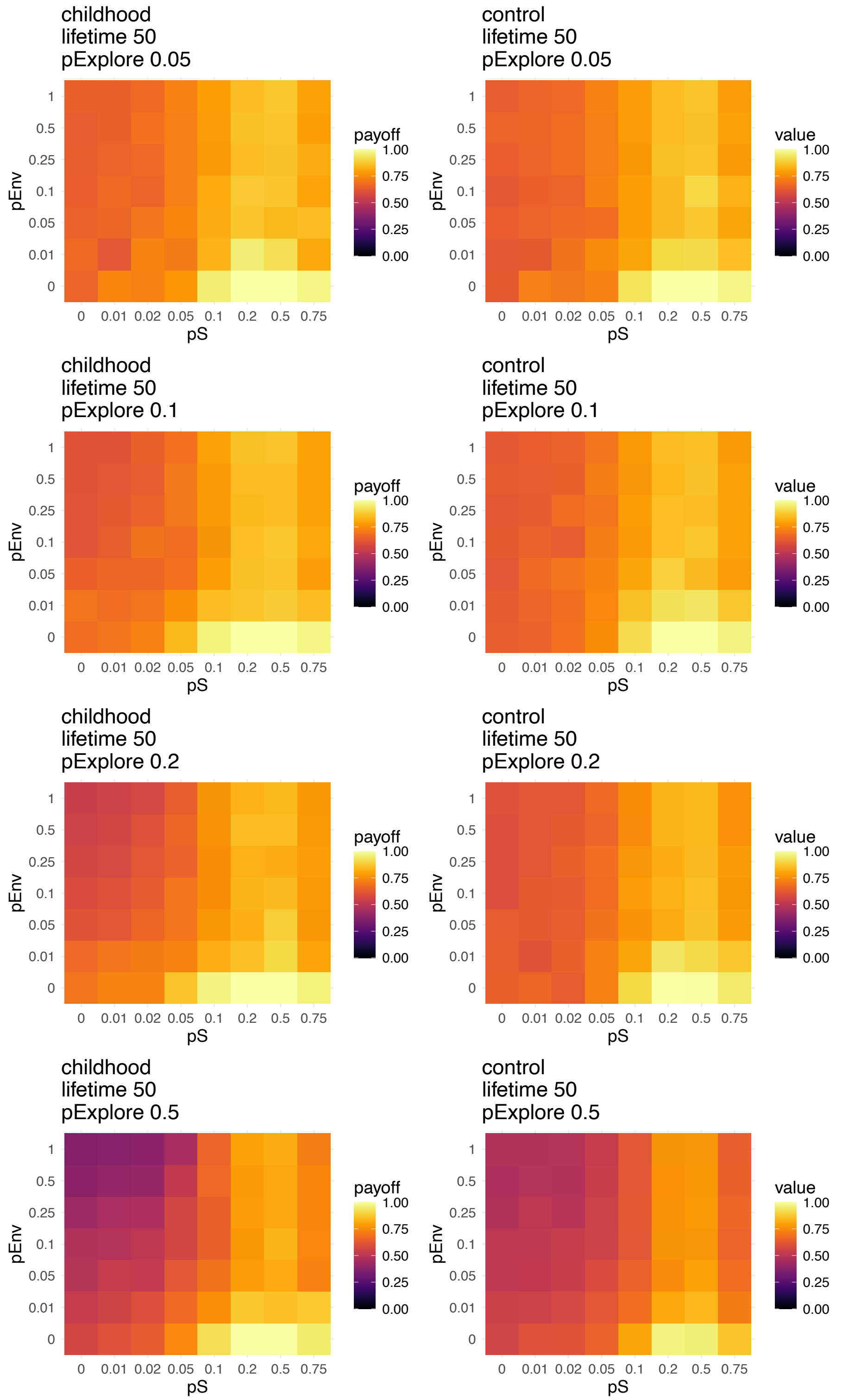


**Fig. S15** – Median values of maximum payoffs varying probabilities of social learning *pS* (when social learning was **random**) and varying probabilities of environmental change *pEnv.* Medians calculated over 100 agents, 20 repeat simulations, and over the last generation of a 5-generation run, and *lifetime* = 50, for all values of *pExplore*.

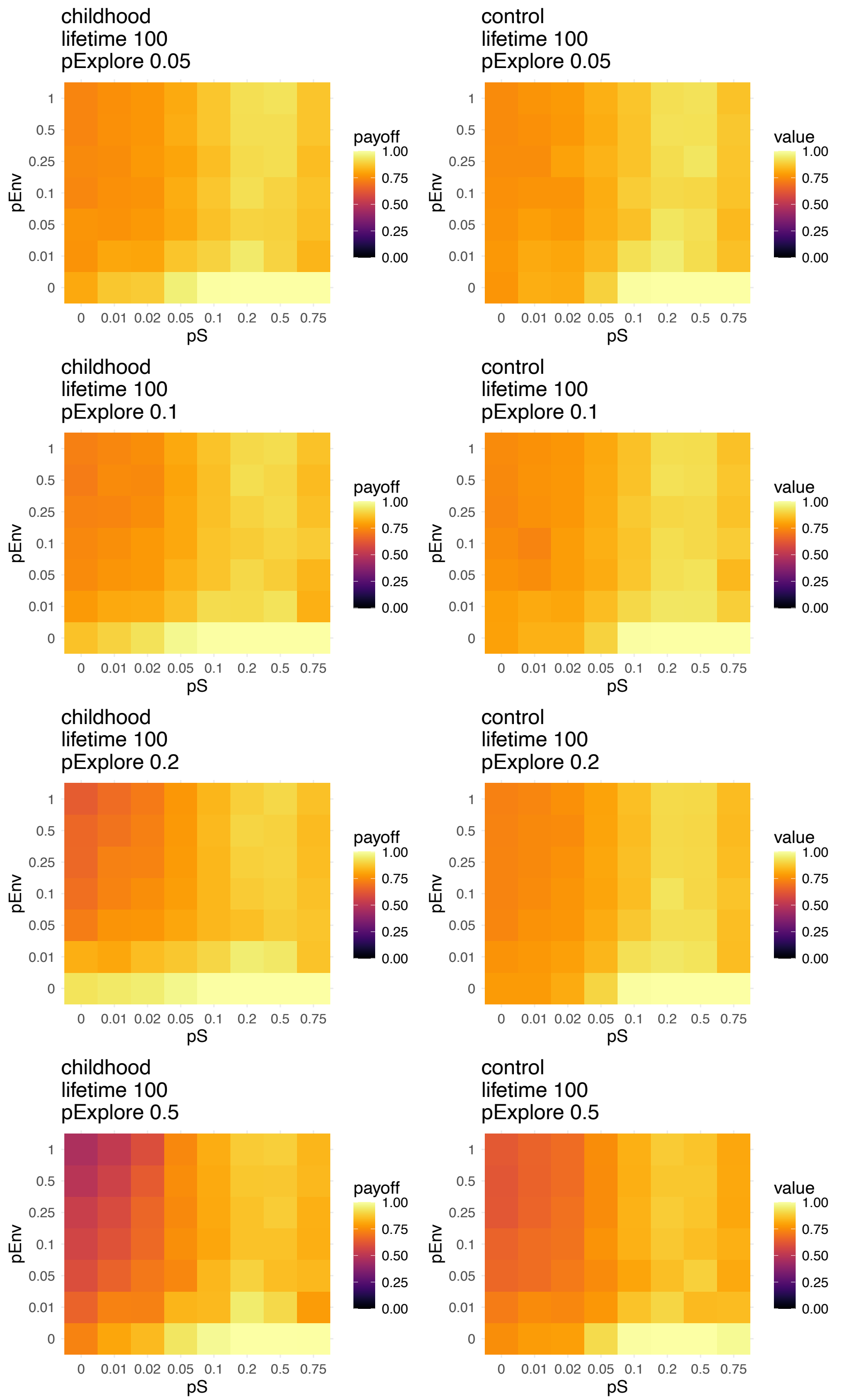


**Fig. S16** – Median values of maximum payoffs for varying probabilities of social learning *pS* (when social learning was **random**) and varying probabilities of environmental change *pEnv*. Medians calculated over 100 agents, 20 repeat simulations, and over the last generation of a 5-generation run, and *lifetime* = 100, for all values of *pExplore*.

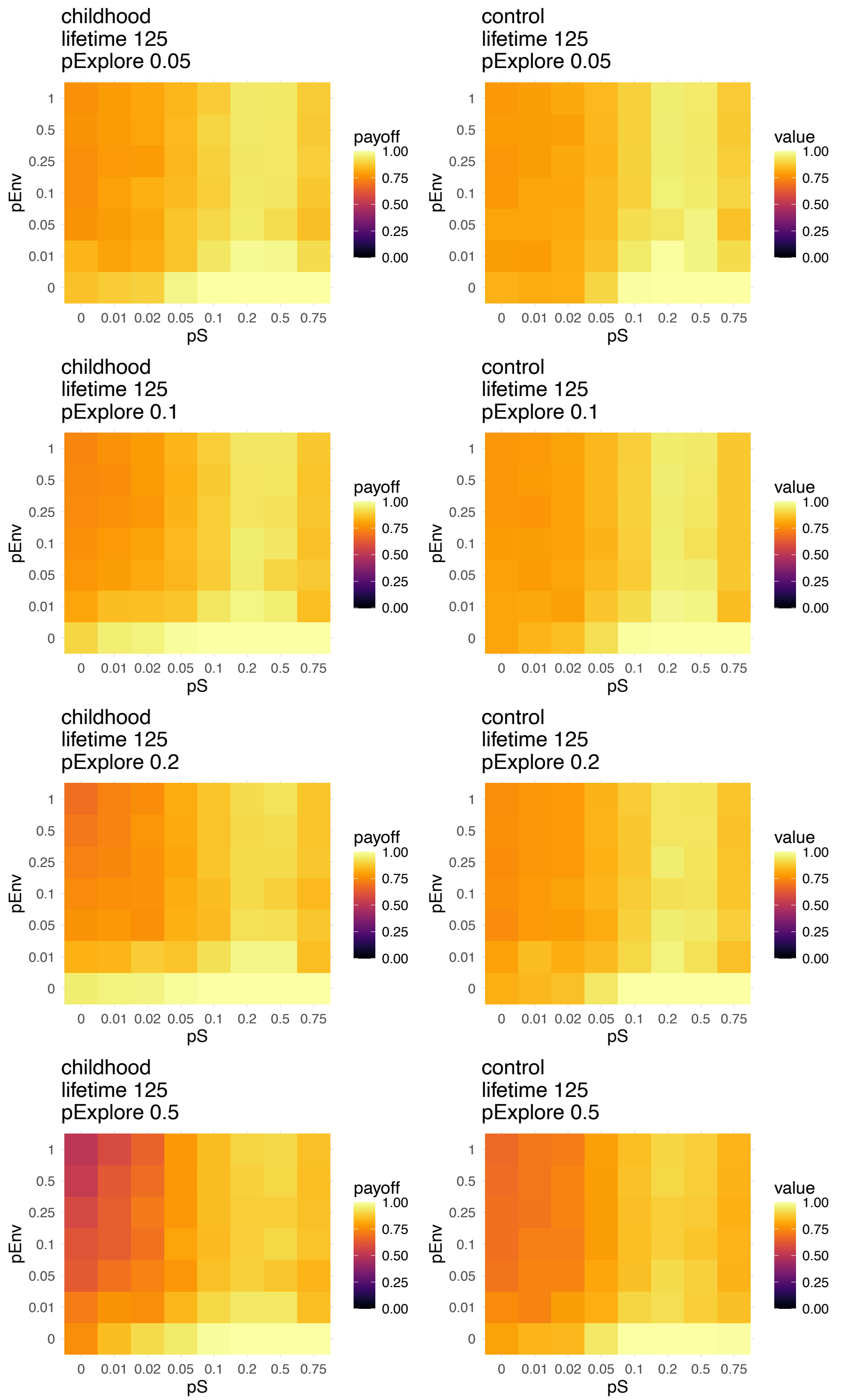


**Fig. S17** – Median values of maximum payoffs for varying probabilities of social learning *pS* (when social learning was **random**) and varying probabilities of environmental change *pEnv.* Medians calculated over 100 agents, 20 repeat simulations, and over the last generation of a 5-generation run, and *lifetime* = 125, for all values of *pExplore*.

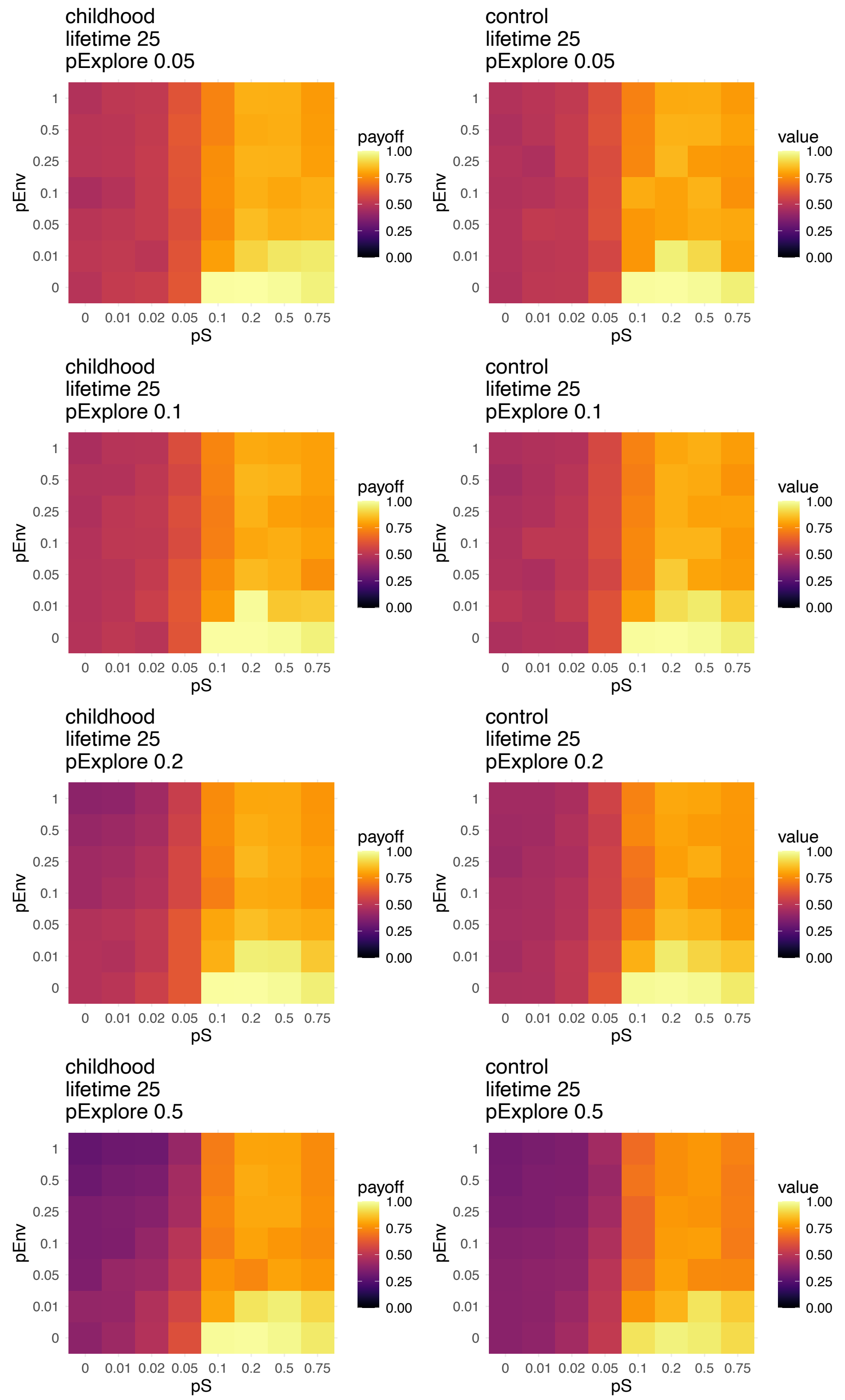


**Fig. S18** – Median values of maximum payoffs for varying probabilities of social learning *pS* (when social learning was **payoff-biased**) and varying probabilities of environmental change *pEnv*. Medians calculated over 100 agents, 20 repeat simulations, and over the last generation of a 5-generation run, and *lifetime* = 25, for all values of *pExplore*.

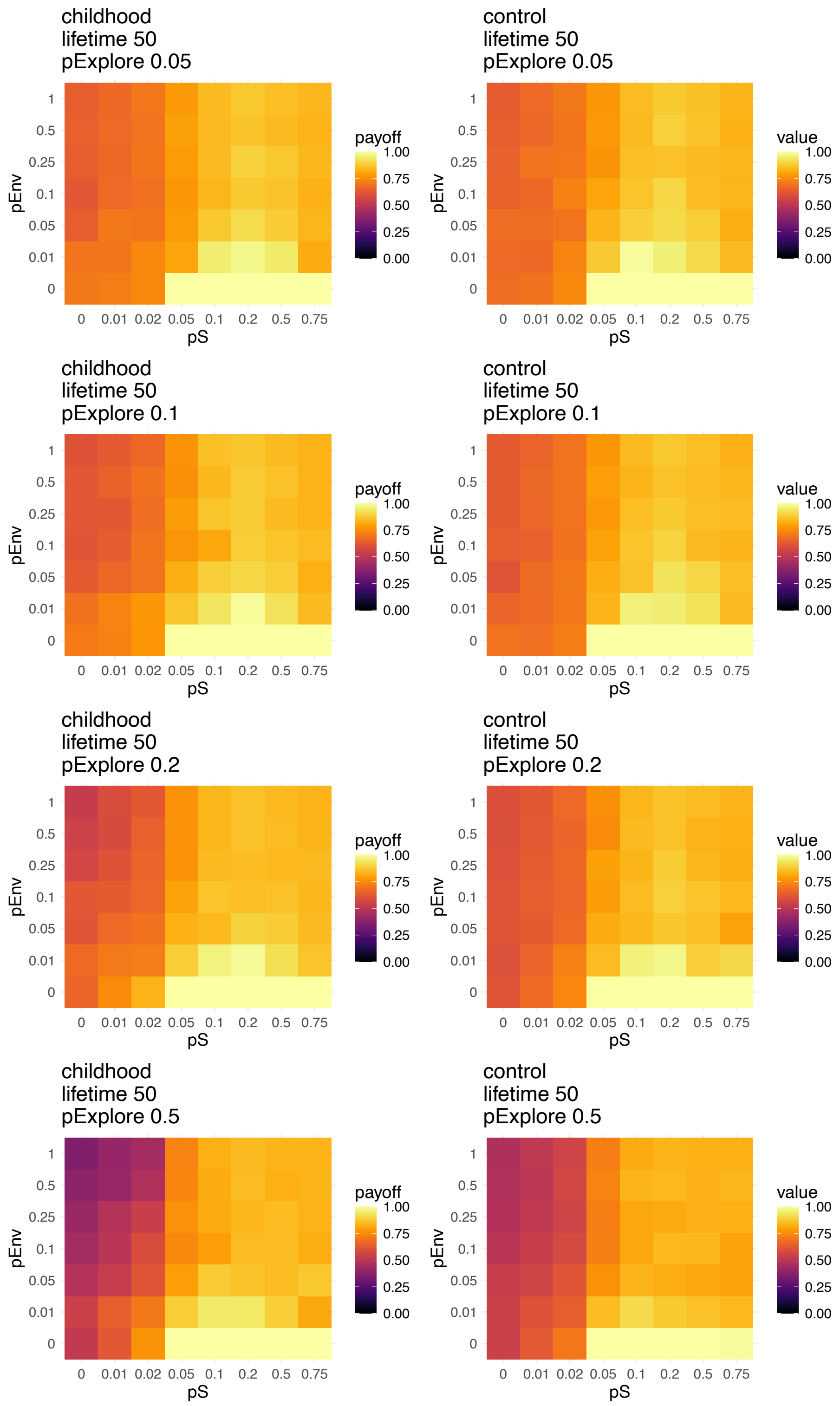


**Fig. S19** – Median values of maximum payoffs for varying probabilities of social learning *pS* (when social learning was **payoff-biased**) and varying probabilities of environmental change *pEnv*. Medians calculated over 100 agents, 20 repeat simulations, and over the last generation of a 5-generation run, and *lifetime* = 50, for all values of *pExplore*.

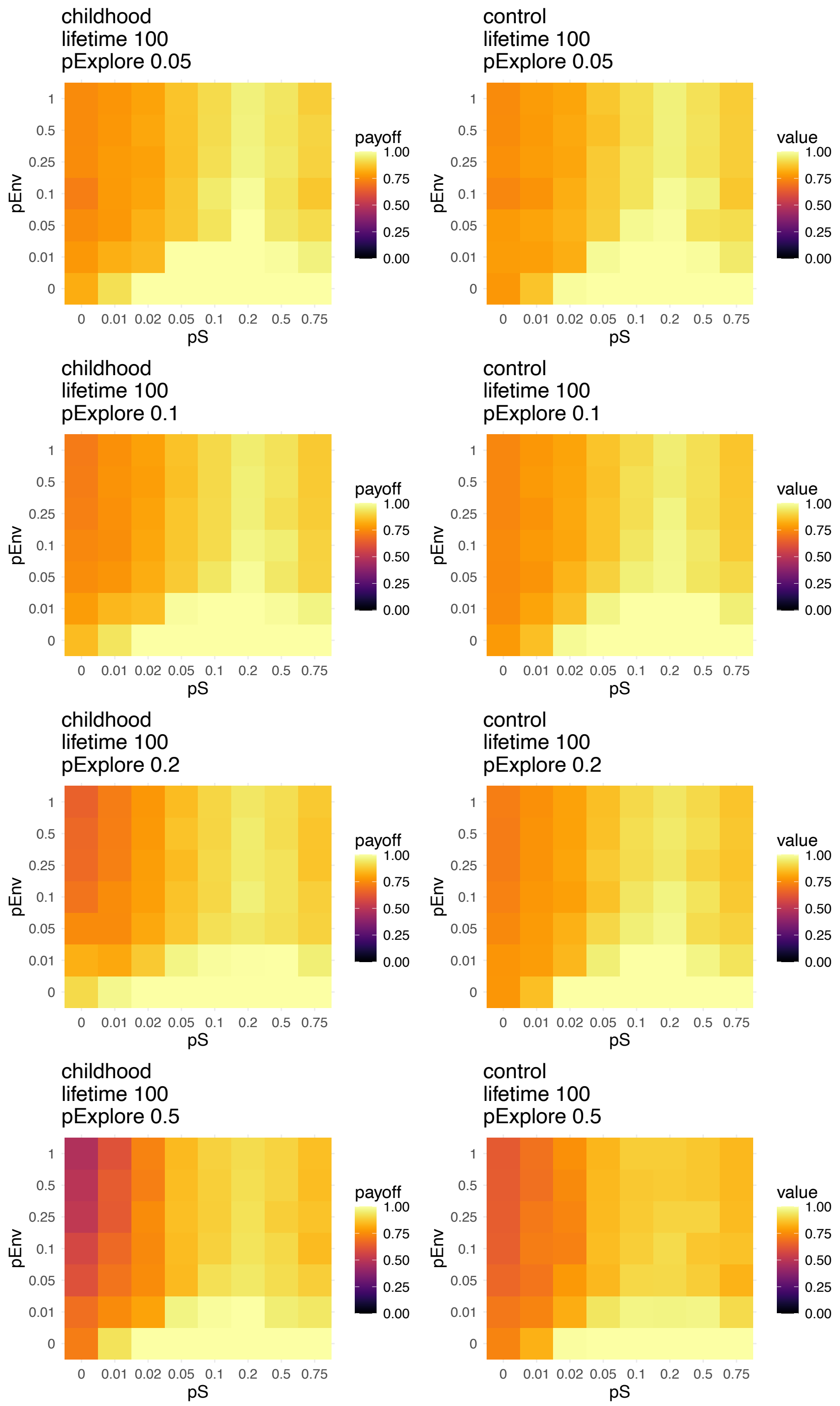


**Fig. S20** – Median values of maximum payoffs for varying probabilities of social learning *pS* (when social learning was **payoff-biased**) and varying probabilities of environmental change *pEnv*. Medians calculated over 100 agents, 20 repeat simulations, and over the last generation of a 5-generation run, and *lifetime* = 100, for all values of *pExplore*.

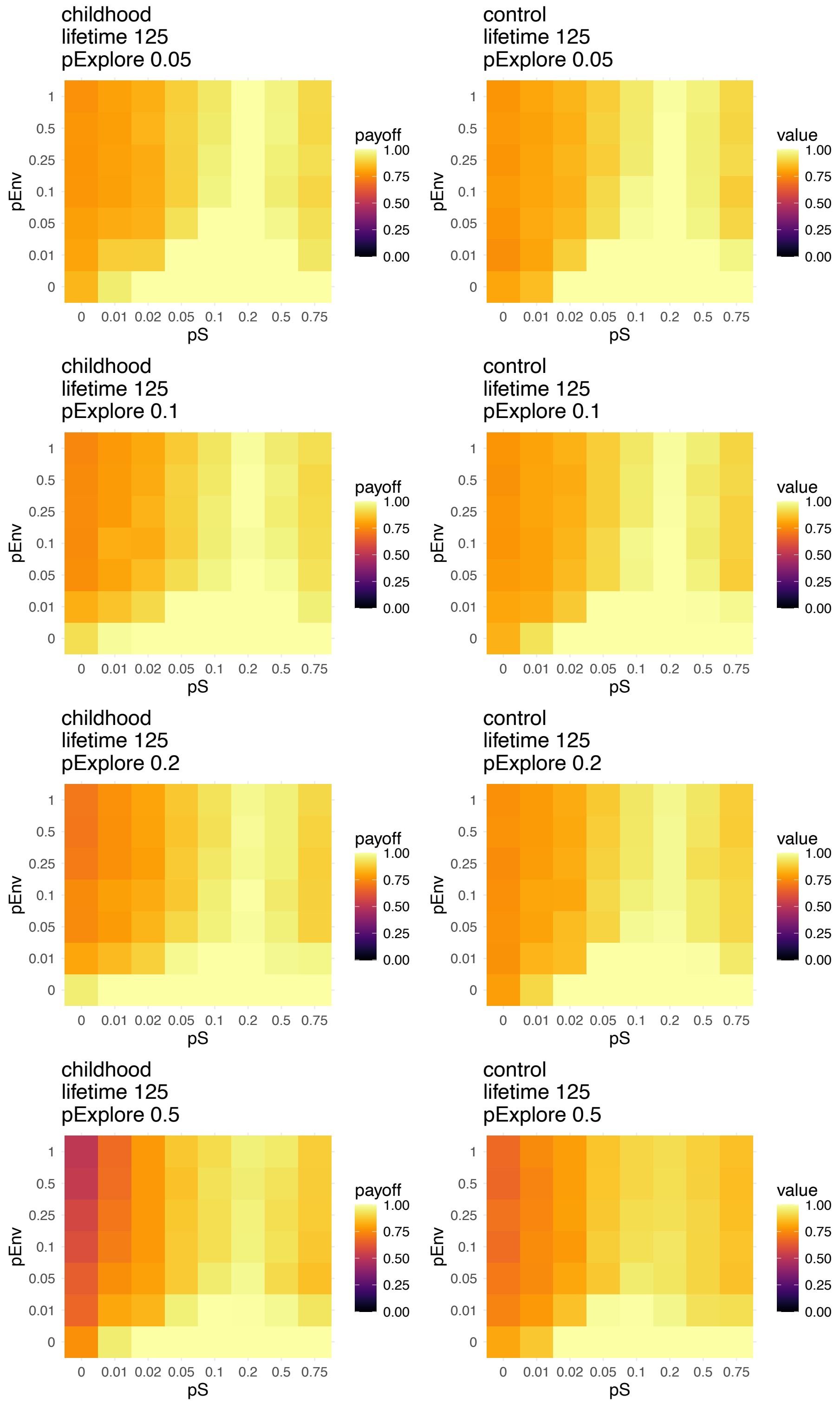


**Fig. S21** – Median values of maximum payoffs for varying probabilities of social learning *pS* (when social learning was **payoff-biased**) and varying probabilities of environmental change *pEnv*. Medians calculated over 100 agents, 20 repeat simulations, and over the last generation of a 5-generation run, and *lifetime* = 125, for all values of *pExplore*.

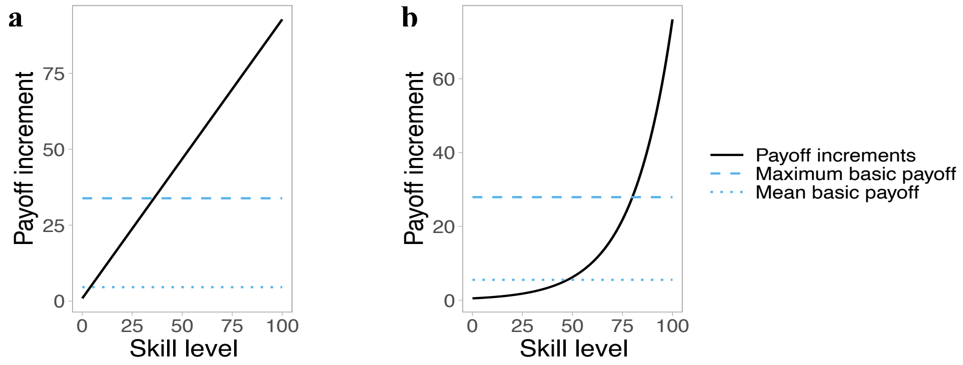


**Fig. S22** - Payoff increment shape and amplitude relative to base payoffs, for (a) linear increments and (b) exponential increments

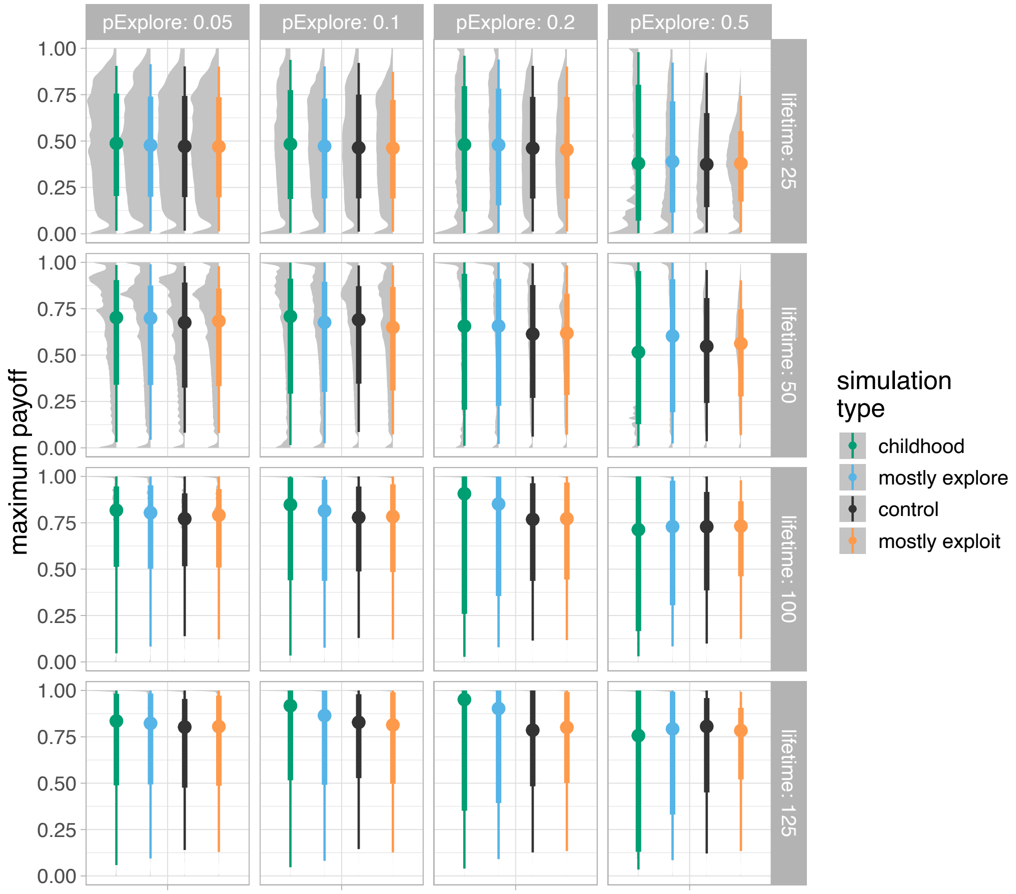


**Fig. S23** - Distributions of maximum payoff for varying lifetime lengths and varying pExplore (i.e. probability agents use explore/proportion of childhood) for all four types of simulations, including additonal intermediate explore conditions. Distributions over 100 agents, 20 repeat simulations, and over the last generation of a 5-generation run. The points indicate the median of the distribution, the thick line indicates the interquartile ranges, and the shape indicates the shape of the density distribution.

### Supplementary text

#### Robustness checks

In order to ensure that our results are not artefacts of the model set-up, we accounted for two critical factors that migth interfere with how populations recover from environmental change. Firstly, when the increment curve had a diminishing returns shape, exploitation very quickly increased payoffs above what the average or maximum payoffs for skill level 1 were. After an environmental change event, therefore, the already exploited behaviours would still be more advantageous than exploiting something new from scratch. This would disadvantage the childhood strategy relative to the control, as its upper hand relied on the initial exploration phase. In order to verify that our results are not purely due to the shape of the increment curve, we ran sets of simulations under two additonal increment shapes: 1. A linear shape, where each increase in level was linearly related to an increase in increment; 2. An exponential shape, where initially improvement was not very useful, but with enough investment improvement became increasingly useful (Figs. S22). We found that the results presented here replicated for both additional shapes under a broad set of parameter combinations (figures not included).

The second factor that could prevent populations from recovering after environmental change more successfully relates to how exploitation was set up in out model. In the results presented here, exploitation involved picking the option with the highest payoff and exploiting it. This, combined with the fact that exploitation results in higher payoff than exploration most of the time, incentivised agents to keep exploiting the same behaviour rather than exploring after the environment changed. In order to allow agents more freedom and an incentive to explore more broadly, we changed the way exploitation took place: rather than picking the best behaviour, the agents would randomly sample *n* behaviours from their repertoire and exploit the best of the chosen *n.* When *n* = *nTraits* this corresponded to the scenario presented here, where agents pick the best of all traits, and when *n*=1 agents exploited a randomly chosen trait from what they knew. We ran simulations with intermediate values of *n*=3, *n*=5, and *n*=10 and. The results varied slightly in terms of which *pExplore* value maximised the basic results without social learning and environmental change – this could be explained by the fact that higher *pExplore* value were now disadvantaged. When *pExplore* was small, agents randomly chose between a small number of traits to exploit, distributing a fixed number of exploit moves to this small subset of options. When *pExplore* was high, they would distribute the same number of exploit moves to a higher subset of options, which resulted in lower exploit levels (and therefore lower payoffs) per trait. However, when we added social learning and environmental change, we found very similar patterns to our main results in terms of the benefits of social learning and the decreases in payoff due to environmental change (figures not included).

Finally, a priori we expected larger differences between the childhood condition and the control condition. To check the robustness of our results, we ran two additional conditions that manipulated the variation in how agents use the explore and exploit moves. In our childhood condition, agents used explore during childhood 100% of the time, and exploit during adulthood 100% of the time. The two new conditions we ran allowed some mixing of explore and exploit during both phases. In the first, we called ‘mostly explore’, agents used explore during childhood

75% of the time, and in the second, ‘mostly exploit’, agents used explore only 25% of the time during childhood. In order to keep the number and proportion of explore and exploit moves consistent across conditions, the proportion was time they used explore during adulthood varied according to their lifetime and the respective length of childhood. On average, these provide intermediate conditions such that in ‘childhood’ agents explored during childhood the most, in ‘mostly explore’ they explore slightly less during childhood, in ‘control’ they explore even less, and in ‘mostly exploit’ they explore the least during childhood. As expected, the childhood condition achieved the highest payoffs, followed by ‘mostly explore’, then ‘control’, then ‘mostly exploit’ (Fig. S23). Under social learning and environmental change conditions the previous results held, with the order mentioned here conserved across all conditions.
